# Comparative genomic analysis identifies potential adaptive variation and virulence factors in *Mycoplasma ovipneumoniae*

**DOI:** 10.1101/2024.05.14.594237

**Authors:** Kimberly R. Andrews, Thomas E. Besser, Thibault Stalder, Eva M. Top, Katherine N. Baker, Matthew W. Fagnan, Daniel D. New, G. Maria Schneider, Alexandra Gal, Rebecca Andrews-Dickert, Samuel S. Hunter, Kimberlee B. Beckmen, Lauren Christensen, Anne Justice-Allen, Denise Konetchy, Chadwick P. Lehman, Kezia Manlove, Hollie Miyasaki, Todd Nordeen, Annette Roug, E. Frances Cassirer

## Abstract

*Mycoplasma ovipneumoniae* is associated with respiratory disease in wild and domestic Caprinae globally, with wide variation in disease outcomes within and between host species. To gain insight into phylogenetic structure and mechanisms of pathogenicity for this bacterial species, we compared *M. ovipneumoniae* genomes for 99 samples from six countries (Australia, Bosnia and Herzegovina, Brazil, China, France, USA) and four host species (domestic sheep, domestic goats, bighorn sheep, caribou). Core genome sequences of *M. ovipneumoniae* assemblies from domestic sheep and goats fell into two well-supported phylogenetic clades that are divergent enough to be considered different bacterial species, consistent with each of these two clades having an evolutionary origin in separate host species. Genome assemblies from bighorn sheep and caribou also fell within these two clades, indicating multiple spillover events, most commonly from domestic sheep. Pangenome analysis indicated a high percentage (91.4%) of accessory genes (i.e., genes found only in a subset of assemblies) compared to core genes (i.e., genes found in all assemblies), potentially indicating a propensity for this pathogen to adapt to within-host conditions. In addition, many genes related to carbon metabolism, which is a virulence factor for Mycoplasmas, showed evidence for homologous recombination, a potential signature of adaptation. The presence or absence of annotated genes was very similar between sheep and goat clades, with only two annotated genes significantly clade-associated. However, three *M. ovipneumoniae* genome assemblies from asymptomatic caribou in Alaska formed a highly divergent subclade within the sheep clade that lacked 23 annotated genes compared to other assemblies, and many of these genes had functions related to carbon metabolism. Overall our results provide evidence that adaptation of *M. ovipneumoniae* has involved evolution of carbon metabolism pathways and virulence mechanisms related to those pathways. The genes involved in these pathways, along with other genes identified as potentially involved in virulence in this study, are potential targets for future investigation into a possible genomic basis for the high variation observed in disease outcomes within and between wild and domestic host species.

**Data Summary:** Raw sequence data and genome assemblies generated for this study have been deposited with the National Center for Biotechnology Information (NCBI) under BioProject number PRJNA1070810. Assemblies are also currently available for download through Dryad with the following link: https://datadryad.org/stash/share/aNet7o-xag3PTjJ0_A_BDoOPUpHHshArGW1eJMfLYl4

NCBI accession numbers and associated metadata for each assembly are available in the Supplemental Materials. DNA sequences extracted from these assemblies for four genetic markers (gyrB, rpoB, 16S, IGS) are available in the Supplemental Materials. Analysis code is available at https://github.com/kimandrews/Movi and an interactive phylogeny is available at https://nextstrain.org/community/narratives/kimandrews/Movi

**Impact statement:** *Mycoplasma ovipneumoniae* causes respiratory disease in wild and domestic sheep and goats around the world, resulting in economic losses for the domestic sheep industry and severe population declines for wild species. Disease outcomes vary widely within and between host species, and this variation could be influenced by genomic differences across bacterial strains. We compared *M. ovipneumoniae* genomes from six countries and four host species and found species-level divergence for strains from domestic goats versus domestic sheep, indicating separate evolutionary origins in these two host species. All wildlife strains fell within these two groups, providing evidence that these strains originated by transmission from domestic populations. We identified genes potentially involved in adaptation to hosts, which could be responsible for differences in disease outcomes across bacterial strains and host species. Many of these genes had functions related to carbon metabolism, a potential virulence factor for Mycoplasmas.

## Introduction

*Mycoplasma ovipneumoniae* (also referred to as *Mesomycoplasma ovipneumoniae* [1, 2]) is a common cause of respiratory disease in domestic sheep (*Ovis aries*) and goats (*Capra hircus*) across the globe, and may be increasing in prevalence in many countries [3-5]. Although *M. ovipneumoniae* infection does not typically cause high mortality rates in domestic flocks, it can significantly decrease lamb production, resulting in a cumulative economic burden [6-9]. This pathogen has also been reported in wild Caprinae species and occasionally other free-ranging and captive ungulates, likely resulting from spillover from domestic sheep and goats [5, 10-12]. In contrast to the relatively low mortality rates caused by *M. ovipneumoniae* in domestic species, infection with this pathogen has been associated with severe pneumoniae in wildlife species including many populations of bighorn sheep (*Ovis canadensis*), free-ranging Norwegian muskox (*Ovibos moschatus*), and a population of captive Dall’s sheep (*Ovis dalli dalli*) [12-15]. Furthermore, *M. ovipneumoniae* infection can impose long-term constraints on population growth in wild bighorn sheep populations, likely due to persistent transmission from chronic adult carriers to neonates [15-18]. However, disease outcomes can vary widely both within and between host species [19-22], and the factors driving this variation remain unknown. Genetically based differences in pathogenicity mechanisms across strains may be a factor contributing to this variation. Mechanisms of pathogenicity are poorly understood for *M. ovipneumoniae*, but several virulence factors have been proposed, including the presence of a polysaccharide capsule, production of hydrogen peroxide as a by-product of metabolism of the carbon-containing molecule glycerol, secretion of hemolysins, and invasion of host cells to evade the host immune response [23-27].

Comparative genomics is a powerful tool that can help uncover mechanisms of pathogenicity and identify potential gene targets for assessing the role of genomic variation in disease outcomes. This approach involves sequencing entire bacterial genomes and then comparing genomic composition across strains and species. For bacterial species with unknown virulence mechanisms, this approach can be used to determine whether genes are present that are known to be virulence factors in other, more well-studied species (e.g., [23, 28, 29]). This approach can also be used to identify genomic variation potentially responsible for differences in pathogenicity observed across closely related strains or species. For example, comparison of the genomes of pathogenic versus nonpathogenic strains has been used to identify genes that may be virulence factors for *Mycoplasma* species infecting domestic pigs, crocodilians, and birds [30-33]. Similarly, analyses comparing the genomes of closely related strains across host species, host tissue types, or environments can be used to identify genes, including virulence factors, that may be involved in adaptation to host or environmental factors [34-36].

For *M. ovipneumoniae*, comparative genomic analyses across strains have been limited in part by the scarcity of publicly available genome sequences. However, studies using sequence data from a small subset of the genome (i.e., fewer than ten genetic loci through multilocus sequence typing or “MLST”) have identified two separate, highly divergent *M. ovipneumoniae* clades that are strongly associated with domestic sheep or domestic goat host species, hereafter referred to as the “sheep clade” and “goat clade” [3, 11, 37]. Genetic diversity is high both between and within these two clades [3, 4, 11, 37, 38], and disease outcomes vary widely across populations and host species for both clades [19-22]. Several studies have found evidence that sheep clade strains cause more severe pneumonia in bighorn sheep than goat clade strains [39-42]. In contrast, a sheep clade strain detected across multiple populations of barren-ground caribou (*Rangifer tarandus granti*) and Dall’s sheep in Alaska has not been associated with population-wide mortality events [43]. This variation in disease outcomes could be driven by strain-specific differences in pathogenicity mechanisms that evolved due to adaptive differences across host species, climates, or other environmental variables. Alternatively, variation in disease outcomes could be driven by other factors such as host resistance, coinfections, or environmental variation.

To gain insight into the genomic basis of pathogenicity in *M. ovipneumoniae*, we generated and compared whole genome sequences of this bacterium across multiple geographic regions, host species, and clades. We used several approaches to identify genes that may be involved in adaptation to within-host conditions or habitats, which could be responsible for differences in pathogenicity across strains and host species. First, we identified genes showing evidence for homologous recombination, positive selection, or diversifying selection, each of which could be indicative of adaptive evolution (e.g., [44-47]). We also identified genes for which presence or absence was significantly associated with host species, which could indicate host adaptation. Identification of these genes provides insight into potential mechanisms of pathogenicity for *M. ovipneumoniae*, including mechanisms that may underly differences in disease outcomes within and between host species. Finally, to evaluate whether MLST markers can be used as indicators of genome-wide variation for *M. ovipneumoniae*, we compared divergence estimates and phylogenetic relationships generated using whole genome sequences versus those generated using sequence data from four MLST markers commonly used for genetic monitoring of this pathogen.

## Methods

### Sample collection

Deep nasal swab samples were collected from domestic sheep, domestic goats, bighorn sheep, and caribou across 10 USA states (Fig. 1, Table 1, Table S1). After collection, swabs were immediately transferred to cryotubes containing a preservation medium composed of either sterile PBS with 20% v/v glycerol, or a growth medium such as TSB with at least 15% v/v glycerol. When necessary, these cryotubes were stored at -20°C or -80°C until the enrichment procedure. Samples collected in the field were shipped on ice or dry ice. Samples were collected by state wildlife management agencies or universities in accordance with animal care guidelines and protocols, or were collected under University of Idaho Institutional Animal Care and Use Committee Protocol IACUC-2019-69.

**Figure 1.**
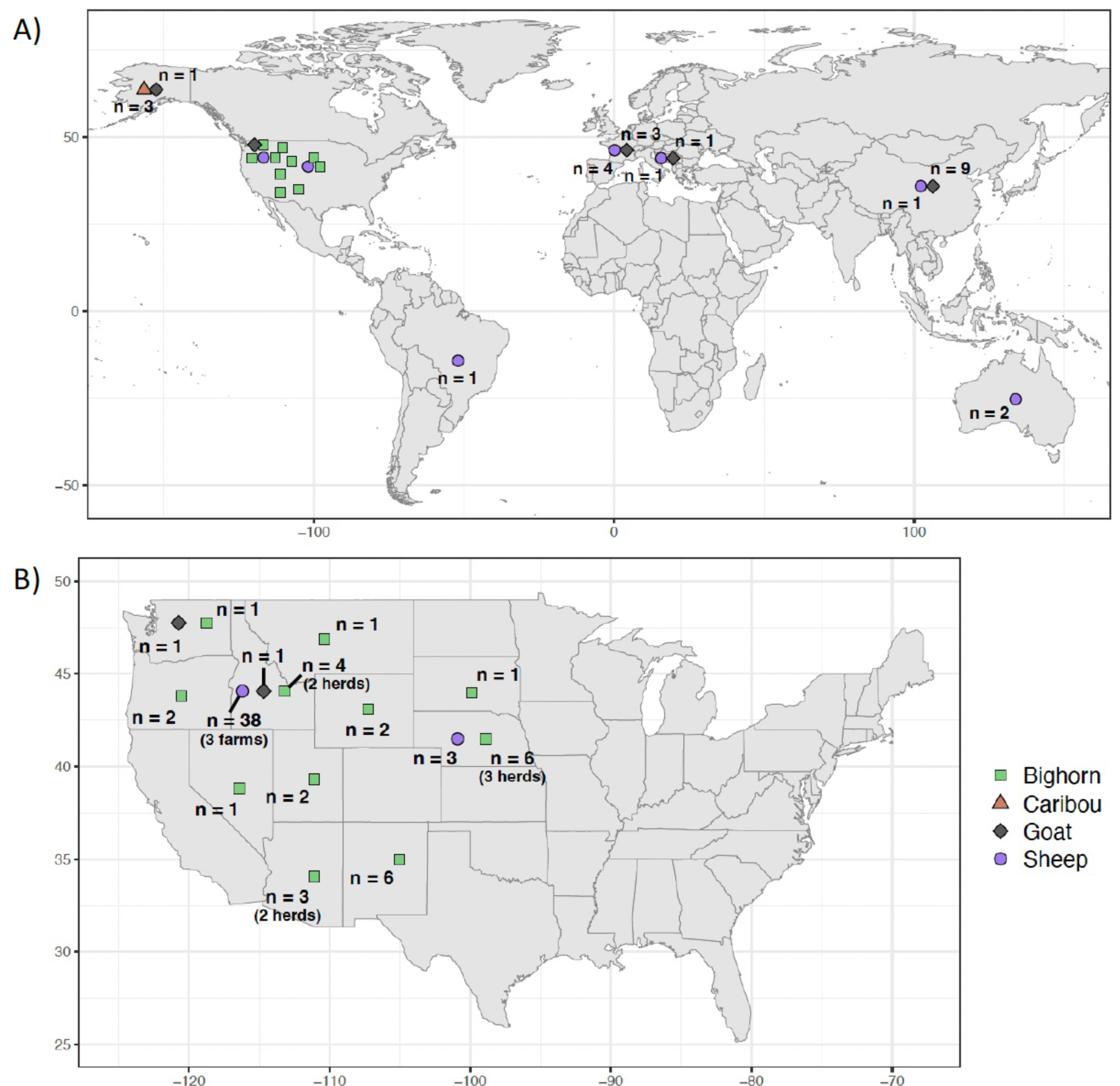
Map of sampling locations for each host species across A) the world; B) 48 USA states. For USA states, the number of herds or farms is shown if more than one was sampled. Locations are shown as the center of the country or state in which the sample was collected, rather than exact sampling location coordinates. The sampling location for the captive bighorn sheep is reported as Nevada; this sheep was infected in South Dakota with a Nevada-origin bighorn sheep strain.

**Table 1.**
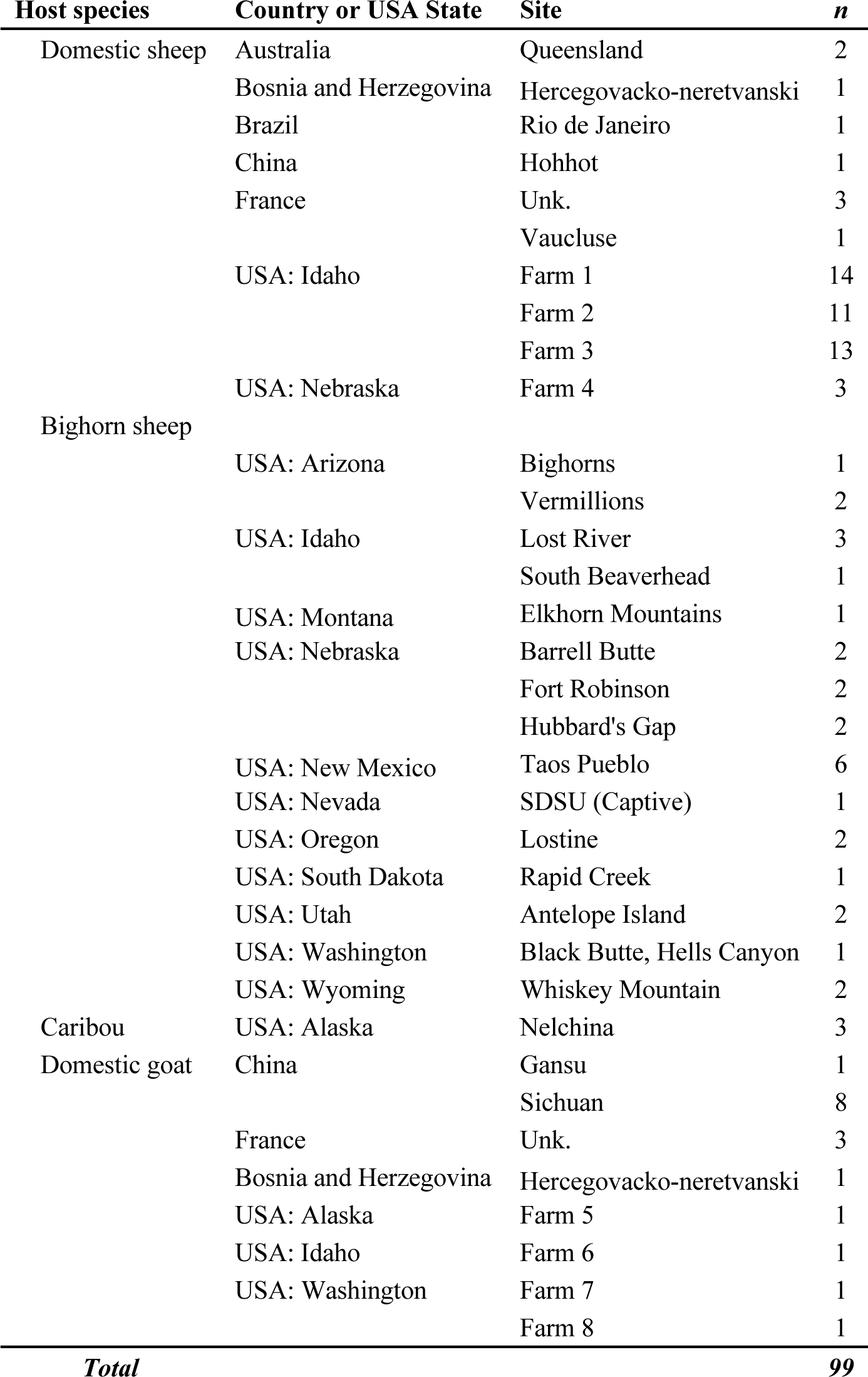
Total sample sizes of *Mycoplasma ovipneumoniae* for each host species and sampling location. “SDSU (Captive)” = captive bighorn sheep that was infected at South Dakota State University with a Nevada-origin bighorn sheep strain. Unk. = unknown.

We also downloaded genome assemblies or raw Illumina shotgun sequence data from NCBI for 25 strains collected from domestic sheep and goats and bighorn sheep from six countries (Australia, Bosnia and Herzegovina, Brazil, China, France, USA; Table S1). These included two assemblies from the type strains of *M. ovipneumoniae* obtained from ATCC (product number 29149, strain NCTC 10151 [Y98]). These strains were originally isolated from domestic sheep in Australia in 1971 [48, 49] (Table S1). For strains that only had raw Illumina shotgun data on NCBI (n=6 from France [3]; Table S1), we assembled the genomes using the method described below for Illumina data. One additional assembly from a domestic sheep in New Zealand was also downloaded from NCBI (GenBank GCA_009756935.1), but this assembly was not included in final analyses due to poor quality (i.e., low completeness and high contamination based on assessment methods described further below).

### Sample culture and enrichment

To selectively enrich *M. ovipneumoniae*, inoculated nasal swabs were subjected to a two-step procedure. First, swabs were inoculated in 4mL mycoplasma broth (product #R102, Hardy Diagnostics, CA, USA) and incubated at 37°C in a shaking incubator (200 rpm) for 4 days. Then, 100 µL aliquots of these enrichment cultures were each transferred into 2 mL of SP4 broth with glucose (Product #U86, Hardy Diagnostics, CA, USA) and re-incubated under the same conditions until either the pH indicator in the culture medium turned orange/yellow (indicating potential growth of glucose-fermenting *Mycoplasma*), or until 15 days had elapsed with no color change observed. Turbid broth cultures indicating the growth of contaminant bacteria, as well as cultures that never turned color, were discarded. After SP4 broth incubation, 1 mL aliquots were removed and centrifuged (20 minutes at 14,000g), and resulting cell pellets were stored at -20°C. For the two samples collected from Wyoming in 2021, a separate protocol was used for culturing *M. ovipneumoniae*, as described elsewhere [50].

### DNA extraction and sequencing

Total genomic DNA was extracted from the frozen cell pellets using the QIAamp DNA Mini Kit (Qiagen). Confirmation of the presence of *M. ovipneumoniae* was obtained by quantitative PCR (qPCR) targeting the 16S rRNA gene using the primers 226-F_new and LMR1, and the 253-P probe, using protocols previously described [6].

Whole-genome shotgun sequencing was performed for all samples that had a qPCR Ct value below 30 (when the threshold was manually set at 0.02). Whole genome shotgun libraries were prepared using Illumina Nextera DNA kits and sequenced on an Illumina MiSeq using the v3 600 cycle sequencing kit. Oxford Nanopore Technologies (ONT) sequencing was also performed for samples with sufficient genomic DNA remaining after Illumina sequencing, and for which analysis of Illumina sequencing data indicated relatively low contamination levels (i.e., the majority of sequence reads were classified as *Mycoplasma*; this analysis is described further below). Barcoded ONT libraries were created using the SQK-LSK109 library prep kit, and sequencing was performed for 48 hours on FLO-MIN106D flowcells, with 12 samples per flowcell. All sequencing was performed at the University of Idaho Institute for Interdisciplinary Data Sciences Genomics and Bioinformatics Resources Core.

### Genome assembly

Raw Illumina sequence reads were cleaned using HTStream (https://ibest.github.io/HTStream) to filter PhiX reads, PCR duplicates, adapter sequences, and low-quality reads. Because *M. ovipneumoniae* is a fastidious microorganism known to present challenges for enrichment culture from nasal swabs, we tested for potential contamination with non-*Mycoplasma* sequences using the metagenomic classifier Centrifuge v1.0.4 (Kim *et al.* 2016) to classify the cleaned Illumina sequence reads. For subsequent analyses, we only retained samples that were dominated by *Mycoplasma* sequences.

For samples with ONT sequence data, raw ONT sequence reads were demultiplexed by barcode and base-called using Guppy v.4.2.2 (Oxford Nanopore Technologies Ltd.). Assembly was performed using Trycycler v0.4.1 [51], which incorporates methods for identifying and removing potential contamination from assemblies, and assemblies were polished using cleaned Illumina sequence reads with Pilon v.1.23 [52] (Supplementary Text). For samples with only Illumina sequence data and no ONT sequence data, cleaned Illumina reads were assembled de novo using SPAdes v3.13.1 [53] using the parameters --careful and -k 33,77,127 (Supplementary Text).

All assemblies, including those from prior studies, were analyzed with CheckM v1.0.13 [54] to evaluate contamination and completeness. This program determines whether expected lineage-specific genes are present in the assemblies. For subsequent analyses, we only retained genome assemblies with low contamination (≤5%) and high completeness (>98%).

### Pangenome characterization

Genome-wide divergence between each pair of samples was estimated by calculating average nucleotide identity (ANI) for all orthologous genes shared between each pair of genomes using FastANI v.1.32 [55]. A threshold ANI value of 95% is often used as a cutoff for the distinction of separate species [55]. All assemblies, including those from prior studies, were initially annotated using Prokka v.1.14.5 [56] with the “-gcode” parameter set to 4 to enable appropriate translation for *Mycoplasma* species. This program uses Prodigal v.2.6.3 [57] for gene prediction, and then predicts gene products by comparing gene sequences against reference databases using BLAST+ v. 2.9.0 [58] and HMMER v.3.2.1 [59]. We then determined the pangenome (i.e., the full set of genes in the dataset) and determined which genes were present or absent in each sample using Roary v.3.13.0 [60]. For this analysis, we used the PRANK [61] codon-aware alignment option and a minimum percent identity of 92%, rather than the default value of 95%, to account for the relatively high level of pairwise genetic variation across samples identified by the ANI analysis (described further in the Results section). We also used the “-z” option to generate sequence alignments for each gene. We then used the sidekick.py script from the HERO pipeline (Park C, Andam C. HERO Github https://github.com/therealcooperpark/hero) to replace the gene-specific headers in the Roary gene alignments with the sample names.

We also annotated the genomes using EggNOG-mapper v.2.1.9 [62, 63]. This approach can outperform Prokka in the number of genes assigned a gene product, especially for non-model organisms including *M. ovipneumoniae*, due to the use of a more comprehensive database [64, 65]. For this analysis, we used the Diamond search strategy [66] and the associated Diamond reference database provided in EggNOG-mapper. As input for this analysis, we used the Roary output file pan_genome_reference.fa, which includes representative sequences from each gene identified by Prodigal during the gene prediction step of the Prokka analysis.

We also used the Roary output to determine the core and accessory genes for our dataset. Core genes are defined as those present in 99-100% of strains, and soft core genes are those present in 95-99% of strains. Accessory genes are comprised of shell genes and cloud genes, where shell genes are those present in 15-95% of strains, and cloud genes are those present in <15% of strains. We also evaluated the total number of core and accessory genes as a function of sample size using Roary. These analyses allow assessment of whether the sample size in our study is sufficient to characterize the core genome and the pangenome of the species. We initially performed Roary analyses using all samples, and then separately for the samples belonging to each of two highly divergent clades identified by our phylogenetic analyses (“sheep clade” and “goat clade,” described further below).

### Virulence genes from prior studies

To identify potential virulence genes in our assemblies, we first used VFanalyzer [67] to identify genes that have been associated with virulence in prior studies for *Mycoplasma*. This program compares genetic sequences against the virulence factor database (VFDB, http://www.mgc.ac.cn/VFs/) [68]. As input for this analysis, we used the Roary output file pan_genome_reference.fa, after converting nucleotide sequences to amino acid sequences. As described above, this file includes representative sequences from each gene identified by Prodigal. We also used the lists of genes identified for each assembly by Prokka and EggNOG-mapper (described above) to determine whether any additional genes were present that were not identified by VFanalyzer but that have been implicated as potentially involved in virulence for *M. ovipneumoniae* and other *Mycoplasma* species in prior studies [23, 65, 69].

### Phylogenetic analysis

We conducted phylogenetic analyses using a nucleotide alignment of all core genes generated using PRANK [61] in Roary. We generated a maximum likelihood tree with RAxML-NG v.1.0.0 [70] with the GTR+Gamma model by first identifying the best-scoring tree from 50 maximum likelihood trees generated from 25 random and 25 parsimony-based starting trees, and then performing bootstrapping for the best tree using default settings.

### Adaptive variation

We used several approaches to identify genes potentially involved in adaptation. First, we identified genes with high levels of homologous recombination, which could be indicative of adaptive evolution [71-73]. We used fastGEAR [74] to quantify recombination for each core and accessory gene, using the gene sequence alignments generated by Roary. FastGEAR first characterizes population structure by identifying lineages using Bayesian Analysis of Population Structure (BAPS) [75] followed by a Hidden Markov Model approach. Next, fastGEAR identifies recent recombinations (occurring at the tips of the phylogeny) and ancestral recombinations (affecting entire lineages) on a gene-by-gene basis. For this approach, recombination sources could be lineages within the dataset, or unknown lineages not represented in the dataset.

Another approach we used to identify genes potentially involved in adaptation was to investigate core genes experiencing positive or diversifying selection. For this approach, we estimated the ratio of nonsynonymous to synonymous substitutions (*dN*/*dS*) for each pair of genomes for each core gene using the codeml function in PAML v.4.9c [76], with alignments for each core gene extracted from the core genome alignment generated by Roary. Values of *dN*/*dS* > 1 indicate positive or diversifying selection, whereas values of *dN*/*dS* < 1 indicate purifying selection. Genes under positive or diversifying selection are candidates for involvement in adaptive differences across strains. For this analysis, we set the codeml “icode” parameter to 3 to enable appropriate translation for *Mycoplasma* species. After generating pairwise *dN*/*dS* values, we filtered out all pairwise comparisons with *dN*/*dS* = 99 (which designates a value that could not be calculated accurately due to a *dS* value near 0 [77]) and all pairwise comparisons with *dN*=0 and *dS*<0.01 (to avoid comparisons between genome sequences from the same population [78]). In addition, we filtered out all pairwise comparisons for genes with poor nucleotide alignments, which we defined as alignments with ≤70% of nucleotides remaining after gaps (i.e., alignment sites with no nucleotides for one or more samples) were removed by codeml.

The next approach we used for identifying genes potentially involved in adaptation was a “pangenome-wide association study” (pan-GWAS) approach implemented in Scoary [79] to identify genes for which presence or absence is significantly associated with one of the hosts (domestic sheep, domestic goats, bighorn sheep, caribou) or each of the two major clades that were identified in our phylogenetic analysis (i.e., the “sheep clade” and the “goat clade”; described further below). The approach used by Scoary accounts for population stratification, and can have strong statistical power even with relatively small sample sizes [79, 80]. For this analysis, we focused on genes that had been assigned a gene product by EggNOG-mapper or VFanalyzer. We defined a gene as “present” in a genome if at least one gene had been assigned that gene product in the genome. Because Scoary requires binary traits, we conducted three tests comparing genomes from the following pairs of groups: sheep versus goat clades (caribou samples were excluded from this analysis since these assemblies had significantly fewer genes than all other sheep clade assemblies, as described further below), domestic sheep versus bighorn sheep hosts within the sheep clade, and caribou samples versus the rest of the sheep clade. We used a Bonferroni correction value of 0.05 as the significance threshold.

Because two of the genes identified as host-associated by the Scoary analysis were part of the CRISPR-Cas system (described further in the Results section), we also investigated whether each genome assembly contained a functional CRISPR-Cas system by identifying high-confidence CRISPR regions (evidence level = 4) using CRISPRCasFinder v.4.3.2 [81-83]. The presence of a functional CRISPR-Cas system in *Mycoplasma* is expected to be characterized by the presence of CRISPR arrays and the genes *cas1*, *cas2*, and *cas9* [84].

To determine whether the genes identified as potential targets of selection were enriched for certain gene functions, we used gene ontology (GO) enrichment analyses implemented in the R package topGO [85]. These analyses were performed using the list of annotated genes in the pangenome as a reference, including all genes that had been assigned a gene product by EggNOG-mapper. We assigned GO terms to each gene in the pangenome using the UniProtKB database [86], including GO terms related to biological processes, molecular functions, and cellular components. We accepted GO terms that had been assigned to bacteria and that had been manually annotated (Swiss-Prot records). For genes that had no Swiss-Prot records, we used GO terms that had been assigned to bacteria in the computationally-annotated TrEMBL database. Genes for which putative gene products had not been identified by EggNOG-mapper were removed from gene lists prior to these analyses. Significant enrichment of GO terms was determined using Fisher’s Exact test with a significance threshold of p<0.05.

### Comparing pathogen monitoring techniques: MLST versus whole genomes

To compare between-sample pairwise genetic divergence values calculated using MLST sequences versus those calculated using genome assemblies, we first used BLAST+ v.2.9.0 [87] to extract sequences from each assembly for each of four MLST markers commonly used for *M. ovipneumoniae* surveillance. These markers include partial sequences from the 16S-23S intergenic spacer region (IGS), the small ribosomal subunit (16S), and genes encoding RNA polymerase B (rpoB) and gyrase B (gyrB) [11]. We conducted this analysis using a custom reference database we created using representative *M. ovipneumoniae* sequences obtained from a prior study for each MLST marker [11]. We then aligned the extracted sequences for each MLST marker using the Geneious Alignment algorithm in Geneious v.11.1.5 (Auckland, New Zealand). For the IGS marker, a portion of the alignment was removed due to the presence of a large number of indels, which led to poor alignment quality that could cause inaccurate estimates of pairwise divergence. We then used the dist.gene function in the R package ape [88] to calculate ANI between each pair of sequences for each marker, for the four markers concatenated, and for all markers except IGS concatenated (since removal of the IGS indel region could have influenced the results). We then plotted the ANI values calculated using MLST markers versus those calculated using the assemblies as described above. We also generated maximum likelihood phylogenies with concatenated MLSTs using RAxML-NG. For this analysis, we partitioned the sequence data by locus and used the GTR+Gamma model for each partition, and used the same tree searching strategy and bootstrapping methods as described above for the genome assemblies.

## Results

### Sample sizes

After quality filtering of genome assemblies, we retained a total of 99 *M. ovipneumoniae* genome assemblies, including 74 that we sequenced and assembled for this study (Supplementary Text), and 25 that we obtained from NCBI (Fig. 1, Table 1, Table S1). Of these, 50 assemblies originated from samples collected from domestic sheep from two USA states and five additional countries (Australia, Bosnia and Herzegovina, Brazil, China, France), including two assemblies from the *M. ovipneumoniae* type strains that were collected from domestic sheep in Australia. In addition, 17 assemblies originated from domestic goats from three USA states and three additional countries (Bosnia and Herzegovina, China, France), 28 from wild bighorn sheep from nine USA states, one from a bighorn sheep infected in captivity with a Nevada-origin bighorn sheep strain, and 3 from caribou from Alaska (Table 1, Table S1). The *M. ovipneumoniae* type strains had been collected in 1971 and one assembly originated from a sample collected from a domestic goat in China in 1998; all other samples had been collected between 2007 and 2021 (Table S1).

### Pangenome characteristics: Full dataset

The *M. ovipneumoniae* pangenome (i.e., the total number of predicted genes across all assemblies) identified by Roary analysis totaled 4,387 (Table S2). The number of core genes was 298 and the number of soft core genes was 79; together, the core and soft core genes represent only 8.6% of the pangenome. For the accessory genes, the number of shell genes was 643 and the number of cloud genes was 3,367. Evaluation of pangenome content as a function of sample size showed that the number of conserved genes leveled off around ten genomes, indicating that the sample size in our study was sufficient to characterize the core genome (Fig. 2). In contrast, the number of accessory genes continued to increase as more genomes were added, indicating that our dataset does not characterize all the accessory genes for *M. ovipneumoniae* (Fig. 2). The high number of accessory genes relative to core genes could indicate strong ability of this pathogen to adapt to within-host conditions, since accessory genes are thought to facilitate adaptation [89, 90].

**Figure 2.**
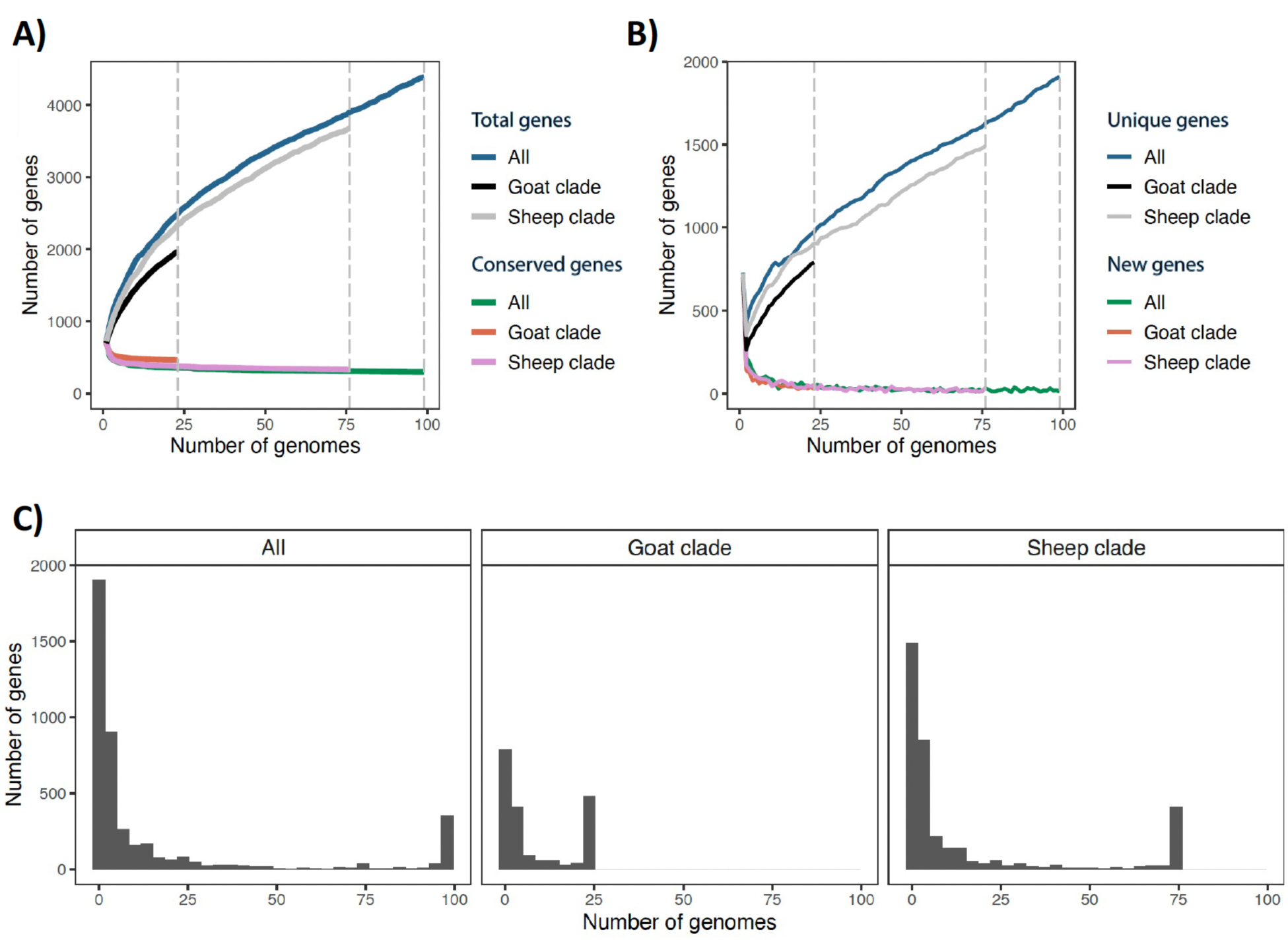
Pangenome characteristics of *Mycoplasma ovipneumoniae* assemblies. Results are shown for all assemblies (n=99), only the assemblies in the sheep clade (n= 76), and only the assemblies in the goat clade (n=23). A) The numbers of genes in the core genome (“conserved genes”) and the pangenome (“total genes”) as a function of sample size. B) The numbers of “new genes” (genes not found in previously sampled assemblies) and “unique genes” (genes found in only one of the sampled assemblies) as a function of sample size. C) The number of genes shared by different numbers of genomes.

Only 726 of the 4,387 predicted genes in the pangenome (16.5%) were assigned a putative gene product by Prokka, with the remaining genes being assigned the category of “hypothetical protein” (Table S2). In contrast, a much higher percentage of the genes in the core genome were assigned a putative gene product (217 of the 298 core genes, 72.8%). A total of 138 annotated genes were present more than once in the pangenome, with the number of copies ranging from 2 to 55 (mean = 3.43) (Table S2). The presence of genes with multiple copies in the pangenome indicates the presence of either paralogous genes or highly divergent copies of orthologous genes that were split into separate genes by the Roary analysis. When using EggNOG-mapper to annotate the genes, a higher percentage of the predicted pangenome genes were assigned a putative gene product (938 genes, 21.4%) than when using Prokka, including 243 (81.5%) of the core genes (Table S2). A total of 179 genes were present more than once, with the number of copies present ranging from 2 to 47 (mean = 3.77) (Table S2). Since EggNOG-mapper assigned products to more genes than Prokka, we used EggNOG-mapper annotations for subsequent analyses investigating adaptive genomic variation (described further below).

### Potential virulence genes

VFanalyzer identified ten genes that are potential *Mycoplasma* virulence factors in our genome assemblies, including enolase (*eno*), hemolysin (*hlyA*), nuclease (*nuc*), and seven genes involved in adhesion (*lppT*, *p102*, *p216*, *p97*, *pdhB*, *plr/gapA*, *tuf*) (Table 2, Table S2). For five of these genes, one copy was found in all 99 genome assemblies (*eno*, *plr/gapA*, *hlyA*, *pdhB*, *tuf*), and for three genes, at least one copy was found in 92-97 assemblies (*lppT*, *p102*, *nuc*) (Table S2). For the remaining two genes, at least one copy was found in 74 assemblies (*p97*) or 10 assemblies (*p216*) (Table S2). Gene annotation with Prokka and EggNOG-mapper also identified additional genes that have been proposed as virulence factors in previous studies of *M. ovipneumoniae* and other *Mycoplasma* species (Table 2, Table S2). These included genes involved in glycerol catabolism and the associated production of hydrogen peroxide (*glpF, glpK, glpO, glpQ*, *gtsA, gtsB, gtsC*), hemolysis (*acpD*), adhesion (*mgpA*), and inflammatory responses (*atpF*) [23, 65, 69]. Each of these additional genes was present in all 99 of our genome assemblies, with the exception of *glpQ*, which was present in only 34 assemblies.

**Table 2.** Genes identified as potentially involved in virulence for *Mycoplasma ovipneumoniae*. Potential mechanism = proposed virulence mechanism for the gene; Analysis^1^ = analysis method that identified the gene as potentially involved in virulence; Sheep clade, Goat clade, Caribou subclade = percent of assemblies within the clade for which the gene was present (Sheep clade excludes the caribou subclade assemblies).

| Gene | Potential mechanism | Analysis | Goat clade | Sheep clade | Caribou subclade |
| --- | --- | --- | --- | --- | --- |
| eno | enzyme | VFDB | 100 | 100 | 100 |
| hlyA | hemolysin | VFDB | 100 | 100 | 100 |
| nuc | nuclease | VFDB | 95.7 | 98.6 | 100 |
| lppT | adhesion | VFDB, Scoary (Caribou vs. Sheep) | 87.0 | 98.6 | 0 |
| p102 | adhesion | VFDB | 87.0 | 100 | 100 |
| p216 | adhesion | VFDB | 13.0 | 5.5 | 100 |
| p97 | adhesion | VFDB | 47.8 | 82.2 | 100 |
| pdhB | adhesion | VFDB | 100 | 100 | 100 |
| plr/gapA | adhesion | VFDB | 100 | 100 | 100 |
| tuf | adhesion | VFDB | 100 | 100 | 100 |
| glpF | glycerol metabolism | Literature | 100 | 100 | 100 |
| glpK | glycerol metabolism | Literature | 100 | 100 | 100 |
| glpO | glycerol metabolism | Literature | 100 | 100 | 100 |
| glpQ | glycerol metabolism | Literature, Scoary (Goat vs. Sheep) | 100 | 21.9 | 0 |
| gtsA | glycerol metabolism | Literature | 100 | 100 | 100 |
| gtsB | glycerol metabolism | Literature | 100 | 100 | 100 |
| gtsC | glycerol metabolism | Literature | 100 | 100 | 100 |
| acpD | hemolysis | Literature | 100 | 100 | 100 |
| mgpA | adhesion | Literature | 100 | 100 | 100 |
| atpF | inflammatory response | Literature | 100 | 100 | 100 |
| rnhB | H<sub>2</sub>O<sub>2</sub> production | Scoary (Goat vs. Sheep) | 39.1 | 0 | 0 |
| baiH |  | Scoary (Caribou vs. Sheep) | 100 | 100 | 0 |
| gcdH |  | Scoary (Caribou vs. Sheep) | 100 | 100 | 0 |
| lemA |  | Scoary (Caribou vs. Sheep) | 100 | 100 | 0 |
| lplA-2 |  | Scoary (Caribou vs. Sheep) | 100 | 100 | 0 |
| nagA | carbon metabolism | Scoary (Caribou vs. Sheep) | 100 | 100 | 0 |
| nagC | carbon metabolism | Scoary (Caribou vs. Sheep) | 100 | 100 | 0 |
| nagE | carbon metabolism | Scoary (Caribou vs. Sheep) | 100 | 100 | 0 |
| nanA | carbon metabolism | Scoary (Caribou vs. Sheep) | 100 | 100 | 0 |
| nanE | carbon metabolism | Scoary (Caribou vs. Sheep) | 100 | 100 | 0 |
| pgmB |  | Scoary (Caribou vs. Sheep) | 100 | 100 | 0 |
| ruvB |  | Scoary (Caribou vs. Sheep) | 100 | 100 | 0 |
| smf |  | Scoary (Caribou vs. Sheep) | 100 | 100 | 0 |
| tdh |  | Scoary (Caribou vs. Sheep) | 100 | 100 | 0 |
| treP | carbon metabolism | Scoary (Caribou vs. Sheep) | 100 | 100 | 0 |
| ulaA | carbon metabolism | Scoary (Caribou vs. Sheep) | 100 | 100 | 0 |
| cas1 | CRISPR-Cas | Scoary (Caribou vs. Sheep) | 100 | 97.3 | 0 |
| cas2 | CRISPR-Cas | Scoary (Caribou vs. Sheep) | 100 | 97.3 | 0 |
| fruA | carbon metabolism | Scoary (Caribou vs. Sheep) | 100 | 97.3 | 0 |
| malC | carbon metabolism | Scoary (Caribou vs. Sheep) | 100 | 97.3 | 0 |
| hsdR |  | Scoary (Caribou vs. Sheep) | 91.3 | 95.9 | 0 |
| malK | carbon metabolism | Scoary (Caribou vs. Sheep) | 100 | 95.9 | 0 |
| pr2 |  | Scoary (Caribou vs. Sheep) | 100 | 95.9 | 0 |

<sup>1</sup>“VFDB” = gene identified by comparing genome sequences against the virulence factor database (VFDB); “Literature” = gene has been proposed as a virulence factor for *M. ovipneumoniae* in previous studies; “Scoary” = gene for which presence/absence was significantly associated with one of the host species through a comparison of the two host species listed in parentheses.

### Phylogenetic analysis

Phylogenetic analyses of the core genes indicated high genetic diversity across *M. ovipneumoniae* and the presence of two highly divergent clades, which were strongly associated with host species. All assemblies from domestic sheep fell in one clade (“sheep clade”), and all assemblies from domestic goats fell in the other clade (“goat clade”) (Fig. 3). All assemblies from bighorn sheep and caribou fell in the sheep clade, with the exception of six assemblies from a single free-ranging bighorn sheep population in New Mexico, which were in the goat clade.

**Figure 3.**
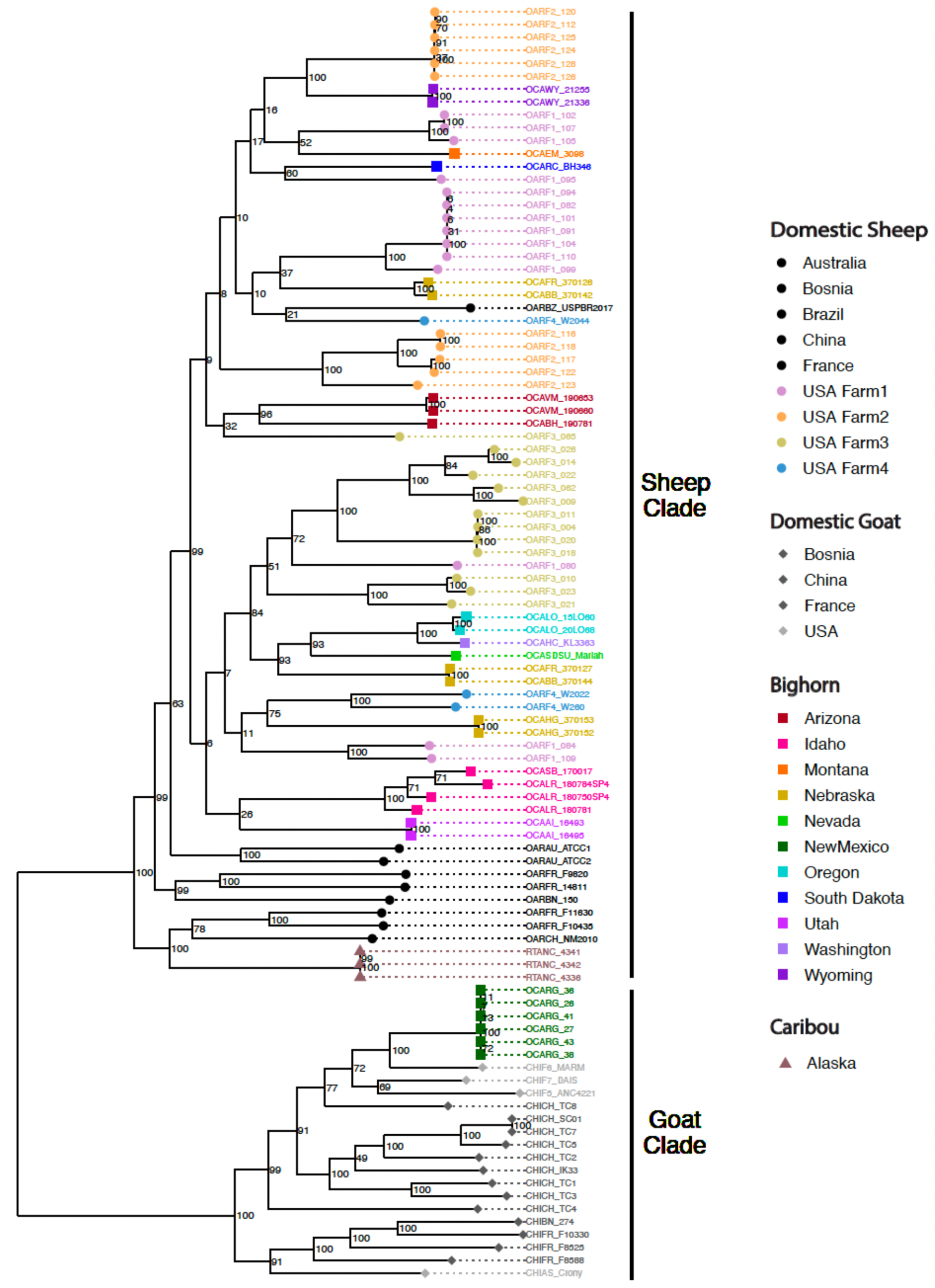
Maximum likelihood phylogeny of core genome sequences for 99 *Mycoplasma ovipneumoniae* assemblies. An interactive phylogeny is available at https://nextstrain.org/community/narratives/kimandrews/Movi

Within the sheep clade, the core genome tree placed all domestic and wild sheep assemblies from USA and Brazil within a subclade with high bootstrap support (value = 99) (Fig. 3). This subclade had minimal phylogenetic clustering associated with geographic distance between herds; for example, each farm had assemblies that clustered in separate, divergent phylogenetic lineages, and some bighorn sheep herds in the same state had highly divergent assemblies, despite their close geographic proximity (e.g., herds OCABB & OCAFR in Nebraska). In addition, USA/Brazil sheep subclade assemblies did not cluster separately by domestic or wild sheep hosts.

The core genome tree indicated the three caribou assemblies were genetically similar to each other and placed these assemblies within the sheep clade, in a separate subclade from the rest of the USA samples, together with two domestic sheep assemblies from France and one from China (Fig. 3). However, the three caribou assemblies were highly divergent from the other assemblies in the subclade, indicating that no close relatives to the caribou assemblies are present in our dataset.

### Genome-wide similarity across assemblies

Estimates of genome-wide similarity across all assemblies indicated the presence of two genetically divergent groups within the full dataset, with ANI values showing a bimodal distribution comprised of one peak around 93% and another around 95.5% (Fig. 4). Within the clades identified by phylogenetic analysis (sheep clade, goat clade, caribou subclade), most ANI values were >94% across pairwise comparisons (Fig. 4). However, all ANI values were <94% for pairwise comparisons between the goat clade and the other groups. These results indicate that the bimodal distribution of ANI values for the full dataset was driven by the strong genetic divergence between the sheep and goat clades. Because a 95% threshold is often used to define separate bacterial species, these results indicate that the sheep and goat clades may represent two separate species.

**Figure 4.**
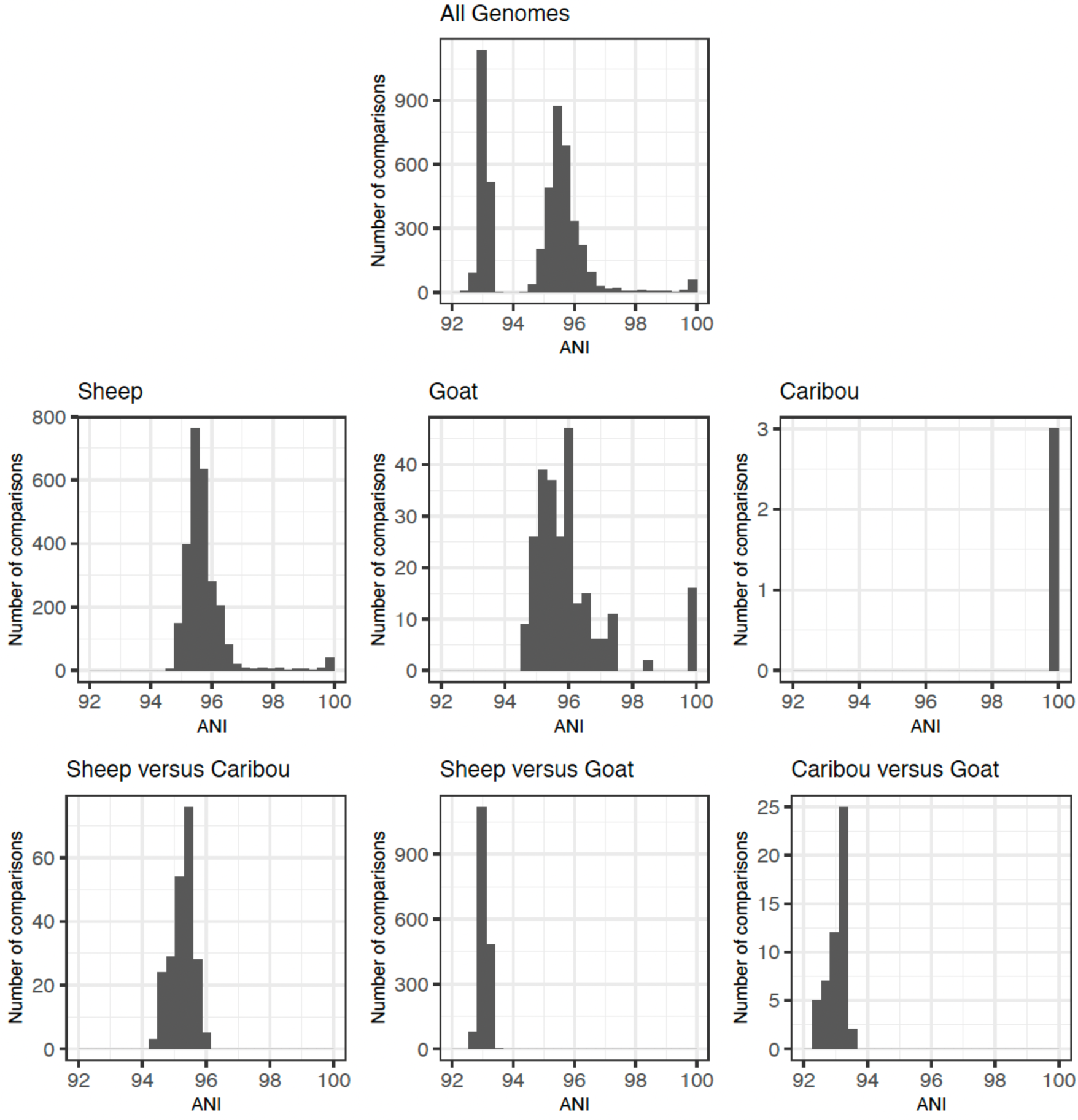
Average nucleotide identity (ANI) for all orthologous genes shared between each pair of 99 *Mycoplasma ovipneumoniae* genomes, including pairwise comparisons across all genomes (“All Genomes”), and pairwise comparisons between and within three groups of samples: “Sheep” clade (including all domestic and bighorn sheep assemblies that were assigned to this clade based on phylogenetic analysis results, but excluding caribou assemblies for comparative purposes); “Goat” clade (including all assemblies assigned to this clade, i.e. all domestic goat assemblies and some bighorn sheep); and “Caribou” subclade (including all assemblies collected from caribou; these formed a subclade within the sheep clade). The greatest genome-wide differences are between the sheep and goat clades; all pairwise comparisons within these clades have ANI >94%, and all pairwise comparisons between these two clades have ANI <94%.

### Pangenome characteristics: Differences between clades and host species

Comparison of assembly lengths across all combinations of clades and host species indicated the three caribou assemblies had smaller genomes than all other samples (ranging from 799,992 bp to 930,395 bp, whereas all other assemblies ranged from 967,712 bp to 1,365,101 bp) (Fig. 5a, Table S1). In addition, the caribou assemblies had fewer genes (mean = 573 genes for caribou, compared to means ranging from 693 to 754 in other clade/host combinations) (Fig. 5b).

**Figure 5.**
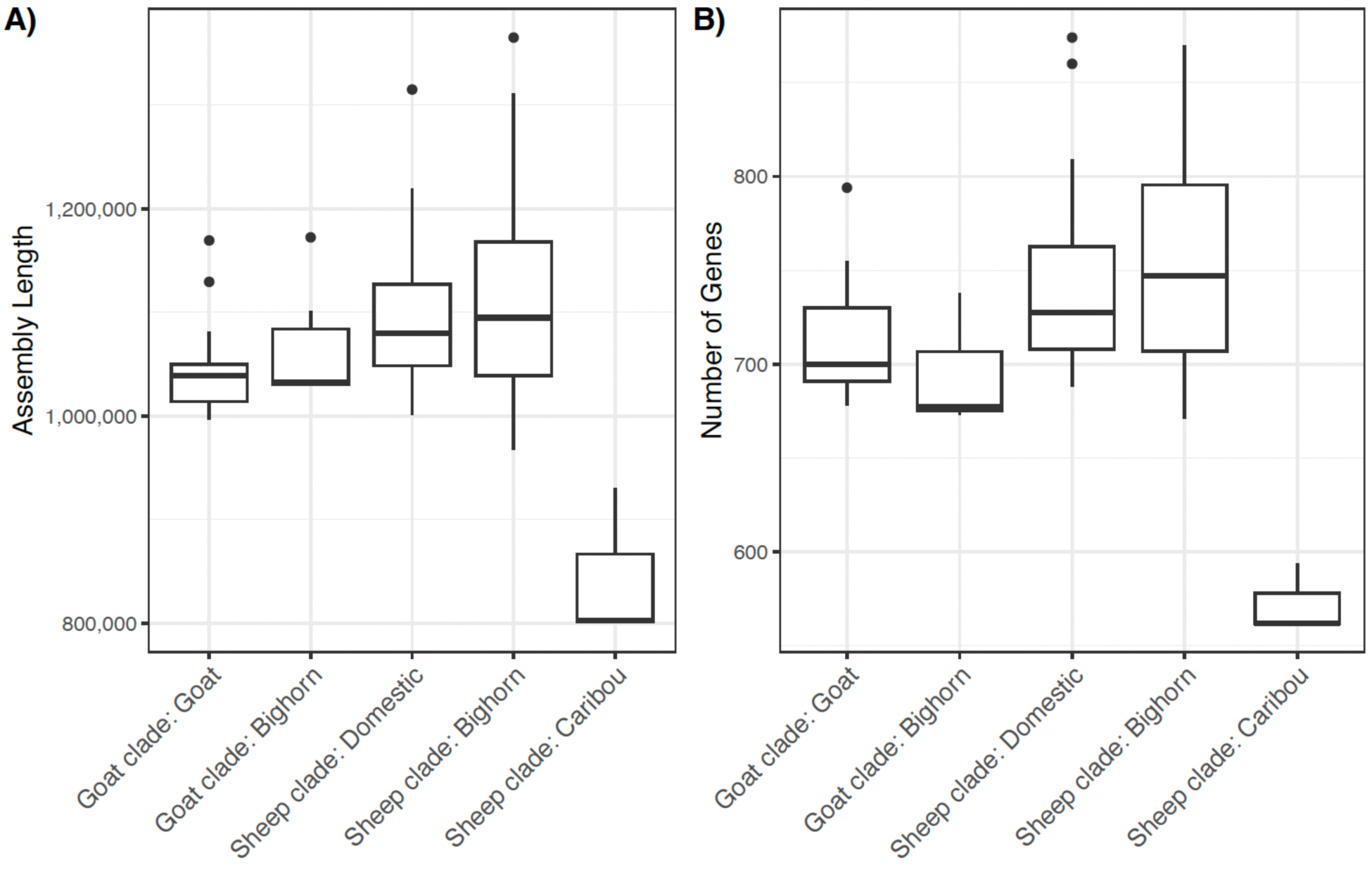
Distributions of a) assembly lengths and b) total numbers of genes for *Mycoplasma ovipneumoniae* genome assemblies across different combinations of clades and host species.

Assembly lengths and numbers of genes were similar across all other combinations of clades and host species (Fig. 5).

Roary analysis with just the goat clade found 464 core genes, 18 soft core genes, 398 shell genes, 1087 cloud genes, and 1967 total genes (Table S3). Roary analysis with just the sheep clade found 331 core genes, 85 soft core genes, 567 shell genes, 2698 cloud genes, and 3681 total genes (Table S4). Rarefaction analyses indicate that the goat clade may have slightly more core genes and fewer accessory genes than the sheep clade (Fig. 2). However, the lower number of genes in the sheep clade is likely influenced by the presence of the caribou assemblies in the sheep clade, given the lower number of genes in caribou assemblies compared to all other assemblies.

### Adaptive variation

*Recombination* The fastGEAR analyses with the full dataset identified 1,069 genes with recent recombinations and 466 genes with ancestral recombinations, of which 384 genes had both recent and ancestral recombinations (Table S5). Of the genes with recent recombinations, 388 were annotated, and 56 of these were present multiple times in this list, indicating paralogs or divergent orthologs in the dataset. When counting only one copy of each annotated gene, a total of 304 unique annotated genes had at least one recent recombination. Of these unique genes, 33 had at least one copy with ≥ 10 recent recombinations. Of the genes with ancestral recombinations, 123 were annotated, of which 20 were present multiple times in the list of annotated genes, and removal of all except one copy of each annotated gene resulted in 96 unique annotated genes. Of these, only one annotated gene (*cas9*) had a copy with ≥10 recombination events; this was the most abundant *cas9* gene copy (present in 70 samples) and had 23 recent recombinations and 20 ancestral recombinations.

Gene ontology enrichment analysis for the annotated genes with at least one recent recombination identified significant GO terms primarily related to transmembrane transport of carbohydrates and other organic substances, providing evidence that recent recombination has been selectively advantageous for genes related to these functions (Table S6). For the set of annotated genes with a large number (≥10) of recent recombinations, GO enrichment analysis identified significant terms related to DNA replication (Table S6). For annotated genes with at least one ancestral recombination, GO enrichment analysis identified significant terms related to the same functions as for genes with recent recombinations (transmembrane transport and DNA replication). This result indicates that homologous recombination may have been selectively advantageous for genes involved in these functions throughout much of the evolutionary history of *M. ovipneumoniae* (Table S6).

*dN/dS* Pairwise comparisons of core genome synonymous and non-synonymous mutations across all samples identified only 22 genes with *dN*/*dS* > 1 (which would indicate positive or diversifying selection) for any pairwise comparisons (Table S7). None of these genes had a large percentage of pairwise comparisons with *dN*/*dS* > 1 (maximum percentage for any gene was 1.5% when considering all comparisons, 3.8% when only comparing goat versus sheep clade genomes, 1.36% when only considering comparisons between sheep clade genomes, and 5.1% when only comparing goat clade genomes; Table S8). Thus, we did not identify any core genes with strong evidence for positive or diversifying selection across our dataset based on *dN/dS* ratios.

*Pan-GWAS* The Scoary analyses with EggNOG-mapper-annotated genes identified two genes that were overrepresented in goat clade assemblies: *glpQ* was present in all goat clade assemblies but only 22% of sheep clade assemblies, and *rnhB* was present in 39% of goat clade assemblies but no sheep clade assemblies (Table 2, Table S9). GlpQ is involved in glycerol catabolism and the associated production of hydrogen peroxide, which is a known virulence factor for *Mycoplasma* [69, 91, 92]. RnhB is involved in DNA repair, and has been shown to protect bacterial cells from oxidative damage in the presence of hydrogen peroxide [93]. Within clades, the presence or absence of these genes did not correspond with phylogenetic relatedness (i.e., within the sheep clade for *glpQ* or within the goat clade for *rnhB*) (Fig. S1, S2). These results indicate that *glpQ* and *rnhB* may be virulence factors associated with the goat clade.

Within the sheep clade, Scoary analysis identified 23 genes that were absent from the caribou assemblies but present in most or all genomes in the rest of the sheep clade, including both domestic and bighorn sheep hosts (Table 2, Table S9). One of these genes (*lppT*) had been identified as a virulence factor by VFanalyzer (described above) due to its involvement in adhesion to host cells for *Mycoplasma* [69]. The remaining genes were enriched for GO terms including catabolic processes, transmembrane transport, and plasma membrane (Table S10, S11). These included multiple genes involved in membrane transport and catabolism of carbon sources alternative to glycerol. The genes absent from the caribou assemblies also included two genes that are part of the CRISPR-Cas system: *cas1* and *cas2* (Table S9, S12). In addition, the caribou assemblies had only partial *cas9* gene sequences (i.e. covering <50% of the consensus sequence from the multi-sample nucleotide alignment) and fewer CRISPR repeats than most other genomes (Supplementary Text, Table S12, S13). These results indicate that the caribou strains do not have a functional CRISPR-Cas system, whereas most other genomes do appear to have functional systems (Supplementary Text, Table S12). No genes were overrepresented in domestic sheep versus bighorn sheep assemblies within the sheep clade.

### Comparing pathogen monitoring techniques

BLAST+ identified one copy of each of the four MLST markers in each assembly, with the exception of two assemblies that lacked a full 16S sequence and two assemblies that had two copies of gyrB (Table S14, Supplemental Results). Pairwise ANI values calculated using MLST sequences were similar to those calculated using whole genomes, ranging from 92.9-100% for within-clade comparisons (compared to 94.4-100% for whole genomes) and 92.4-99.4% for between-clade comparisons (compared to 92.4-93.5% for whole genomes) (Fig. 4, Fig. S3, S4). However, ANI calculated with MLST sequences had greater variance and tended to underestimate divergence compared to ANI calculated with whole genomes, although variance decreased when MLST markers were concatenated (Fig. S3, S4). Phylogenetic analysis with concatenated MLST markers separated the sheep and goat clade with strong bootstrap support (88-94%), but most other internal nodes had low bootstrap support (Fig. S5, S6).

## Discussion

Comparative genomic analysis of 99 *M. ovipneumoniae* genome assemblies from six countries and four host species found evidence for adaptive variation that could be related to differences in pathogenicity mechanisms across strains. We found high genetic diversity and a high proportion of accessory genes compared to core genes in the *M. ovipneumoniae* pangenome, characteristics which are predicted to promote adaptation [89, 90]. We also found that the two major clades that were observed in previous studies to segregate by domestic host species (sheep versus goat) are divergent enough to be considered separate bacterial species, indicating that these clades likely evolved in separate ancestral host species. Within the sheep clade, three closely related assemblies from asymptomatic wild caribou in Alaska had significantly smaller genomes and fewer genes than all other assemblies. These genomic differences indicate a separate evolutionary trajectory for this subclade resulting from geographic isolation or adaptation to a unique host or environment. However, a larger sample size from wildlife populations in Alaska is needed to confirm these results. In contrast, bighorn sheep strains did not cluster separately from domestic sheep strains in phylogenetic analyses, consistent with multiple spillover events from domestic sheep to bighorn sheep rather than separate evolutionary origins in these two hosts [11].

We identified many genes that showed evidence for potential involvement in adaptation based on the detection of homologous recombination, which allows exchange of selectively advantageous genetic variation between bacterial cells [46, 47]. Significant GO terms for these genes were related to the transport of organic substances including carbohydrates, the metabolism of which is a predicted virulence factor in *Mycoplasma* [69]. Furthermore, the significant GO terms included a specific group of transporters, the ATP-binding cassette (ABC) transporters, which are thought to play an important role in virulence for *Mycoplasma* and other bacterial species [94-96]. High homologous recombination in genes encoding transport proteins has also been observed across a wide range of bacterial species [46, 97] and is thought to be driven by the direct contact of transport proteins with the extracellular environment, which could drive strong adaptive responses to environmental change.

We also identified genes for which presence or absence was associated with host species, which are candidate genes for differences in pathogenicity mechanisms resulting from evolutionary origins in separate host species or within-host habitats. Most of the genes identified as associated with host species were involved in carbon metabolism. For example, when comparing sheep versus goat clades, two annotated genes (*glpQ* and *rnhB*) were strongly associated with the goat clade, and both of these have functions related to the production of hydrogen peroxide, which is a byproduct of glycerol metabolism and considered to be a virulence factor for *Mycoplasma* [92]. The gene product of *glpQ*, which was present in all goat clade assemblies but only 22% of sheep clade assemblies, performs enzymatic and regulatory roles in the catabolism of glycerol and its derivative glycerophosphocholine, which result in the production of hydrogen peroxide [69, 91, 92]. The gene product of *rnhB*, which was present in 39% of goat clade assemblies but no sheep clade assemblies, plays a role in DNA repair, and has been shown to protect bacterial cells from oxidative damage in the presence of hydrogen peroxide [93].

When comparing caribou subclade assemblies to the rest of the sheep clade, 23 annotated genes were absent in the caribou assemblies but present in most or all other sheep clade genomes. One of these genes is a potential *Mycoplasma* virulence factor (*lppt*) that plays a role in adhesion to host cells [69]. The remaining 22 genes were enriched for biological processes and molecular functions involving catabolism of carbon sources other than glycerol, including genes encoding membrane transporters and catabolic enzymes, many of which showed evidence for homologous recombination in the analyses described above. These genes were involved in catabolism of sialic acid (*nanA*, *nanE*, *nagA*, *nagC*, *nagE*), N-acetylglucosamine (*nagA*, *nagC*, *nagE*), maltose and maltodextrin (*malC*, *malK*), fructose and mannose (*fruA*), trehalose (*treP*), and ascorbate (*ulaA*). Many of these genes have been proposed as virulence factors in *Mycoplasma* and other bacterial species, including *nanA* and *nanE* for *M. hyorhinis* and *M. alligatoris* [30, 98]; *nagA*, *nagC*, *nagE* for *M. hyopneumoniae* and other bacterial species [30, 99, 100]; *fruA* for *M. gallisepticum* [33, 101]; *treP* for a wide range of bacterial species [102]; and *ulaA* for *M. hyopneumoniae* [103]. The absence of these genes in caribou assemblies suggests that these strains are more restricted in the types of carbon sources that can be utilized compared to other strains. These genes may have been lost over time through reductive evolution if certain carbon sources were absent or redundant in the environment [104] or through adaptive gene loss if the genes were disadvantageous in the environment [105]. The absence of these genes in Alaska wildlife strains could contribute to the apparently low virulence of these strains, potentially due to lower growth opportunities resulting from the inability to use available carbon sources, reduced production of toxic byproducts of carbon catabolism, or other factors. However, a larger sample size of *M. ovipneumoniae* from Alaska wildlife would be needed to confirm that these genes are consistently absent across strains.

The genes absent in caribou assemblies also included *cas1* and *cas2*, which are both critical to the CRISPR-Cas system. Furthermore, other critical components of the CRISPR-Cas system showed evidence for being nonfunctional in these assemblies; the *cas9* gene was truncated, and the number of CRISPR repeats was lower than for most other genomes in our study. In contrast, the CRISPR-Cas system appears to be functional in most other strains in our study. CRISPR-Cas systems provide defense mechanisms for bacteria against foreign invaders such as phages and plasmids, but also play a role in virulence through a wide range of mechanisms [106-108]. For example, the inactivation of *cas9* has been shown to decrease virulence for some bacteria by decreasing adhesion, toxin production, and intracellular survival [107, 109, 110]. Therefore, the absence of a functional CRISPR-Cas system in the caribou strains could be related to the apparently lower virulence of these strains.

Some of our tests regarding adaptation did not result in the identification of candidate genes. For example, we did not identify any genes that were significantly associated with domestic versus bighorn sheep hosts, and therefore we did not identify any genes for which presence or absence was associated with spillover and persistence in bighorn sheep. However, larger sample sizes would be required to fully evaluate whether the presence or absence of certain genes may create an adaptive advantage in bighorn sheep. We also did not identify any core genes with evidence for positive or diversifying selection across our dataset based on *dN/dS* ratios. However, these analyses are only able to identify genes for which selection has caused nonsynonymous nucleotide changes in multiple amino acids, and are unable to identify genes for which selection has acted on only a small portion of the gene, such as a single amino acid. Furthermore, these analyses focused only on core genes, which are predicted to experience strong purifying selection, whereas accessory genes may be more likely to be involved in niche specialization and experience positive or diversifying selection [89].

We also identified ten genes that are present across some or all *M. ovipneumoniae* strains in our dataset that have been implicated as potential virulence factors for *M. ovipneumoniae* and other *Mycoplasma* species in prior studies. These included additional genes involved in adhesion and glycerol metabolism and the associated production of hydrogen peroxide, as well as hemolysis and other functions.

### Phylogeography

We found high genetic diversity for *M. ovipneumoniae* and minimal phylogenetic clustering by geographic distance between herds, consistent with previous MLST studies in the USA and Europe [3, 11, 24]. These patterns likely result from large population sizes and strong anthropogenic determinants of long-distance movement patterns for domestic sheep. However, all USA assemblies, with the exception of the caribou assemblies, clustered into one highly supported clade with a deep ancestral node. This pattern suggests relatively low rates of new introductions of *M. ovipneumoniae* into the USA in recent history. The caribou strains clustered more closely with strains from France and China than from the USA, but nonetheless were highly divergent from those strains, indicating that our dataset does not include any close relatives to the caribou strains. These results suggest that the caribou strains may be descendants of a separate introduction than the introductions from which all other USA strains in our dataset are descended, potentially originating from a geographic region that was not sampled in our dataset. Although our dataset only includes three samples from caribou, the MLST sequences of these three samples are identical to all other *M. ovipneumoniae* from caribou and Dall’s sheep in Alaska sequenced over a 15-year period (2004-2019) [43]. This indicates that *M. ovipneumoniae* in Alaska wildlife populations may be characterized by low genomic diversity, and that the three assemblies in our dataset may be representative for these populations, although more samples would be needed to confirm these patterns. The low MLST diversity of the caribou strains suggests that the introduction from which these strains descended may have been a relatively recent event. However, our analyses cannot identify the geographic source, date, or number of *M. ovipneumoniae* introductions into Alaska wildlife populations; to address these questions, more samples would be needed from more caribou populations and all potential introduction source regions.

### Implications for pathogen monitoring

Genomic monitoring of *M. ovipneumoniae* is typically conducted by sequencing four MLST markers rather than whole genomes, due to the challenges and cost of culturing this pathogen and performing whole genome sequencing. One of the primary uses of MLST sequences has been the tracing of introduction sources for bighorn sheep populations. We found that strains with >99.9% similarity in MLST sequences were also highly similar at a genome-wide level, suggesting that strains with highly similar MLST sequences are likely indicators of recent transmission events. We also found that measures of divergence between strains were overall similar between MLST sequences and whole genome sequences, indicating that MLST sequences can be used as rough estimates of genome-wide divergence. However, MLST divergence estimates had greater variance and tended to underestimate genome-wide divergence, especially between sheep and goat strains. Phylogenies generated with concatenated MLST markers separated the sheep and goat clades with high bootstrap support, indicating that MLST markers can also be used to reliably distinguish these clades.

### Limitations

One limitation of this study was that many of the genes identified in the *M. ovipneumoniae* assemblies remained unannotated due to the current lack of genetic information for this non-model organism. Some of the genes identified as outliers in our analyses were not annotated, and therefore the potential involvement of those genes in pathogenicity remains unexplored. Also, the gene clustering analysis performed by Roary appeared to separate some orthologous genes by clades or subclades due to the high genetic variation in our dataset, causing some orthologous genes that were shared across all assemblies to appear to be present in only one clade. To minimize the separation of orthologous genes, we reduced the Roary minimum percent identity parameter to 92%, with that value chosen based on ANI values (as described in the Methods section); reducing this value further ran the risk of grouping non-orthologous genes. To account for the potential impact of orthologous gene separation on comparisons of gene presence or absence across clades and host species, we used gene annotations rather than gene clusters to define a gene as present or absent in each assembly. Another limitation of this study was the relatively low sample sizes, especially for the caribou strains, for which only three samples were available that had been collected from the same herd within a 24-hour period. MLST analyses indicate very low diversity across Alaska wildlife populations, indicating that these three assemblies may be representative of these populations, but future studies should use more samples to determine whether the patterns observed here are consistent across populations.

## Conclusions

In this study, we substantially increased the number of publicly available genome assemblies for *M. ovipneumoniae*, as well as the geographic and host species ranges of available assemblies, and provided new insight into phylogenetic structure and potential pathogenicity mechanisms for this bacterium. Phylogenetic analyses indicated two clades with species-level divergence that likely evolved separately in domestic sheep versus domestic goat hosts, and high diversity and minimal geographic clustering within those two clades. Comparative genomic analyses identified multiple genes and cellular processes that may be involved in pathogenicity mechanisms across clades and host species. In particular, we found evidence that carbon metabolism may play a role in virulence and adaptation to host immune defenses for *M. ovipneumoniae*. The results of these analyses, along with the database of genome assemblies generated here, provide a resource for identifying genes and cellular processes to target for further investigation into the epidemiology and virulence of this pathogen.

## Ethical approval

Samples were collected under University of Idaho Institutional Animal Care and Use Committee Protocol IACUC-2019-69 or according to approved state wildlife agency animal welfare protocols.

## Author contributions

Conceived study: K.R.A, E.F.C., T.E.B., T.S., S.S.H., E.M.T.; conducted fieldwork and/or provided samples: K.R.A, E.F.C., T.E.B., T.S., K.B.B., L.C., A.J-A., D.K., C.P.L., K.M., H.M., T.N., A.R.; conducted laboratory analyses: T.S., K.N.B., M.W.F., D.D.N, G.M.S, A.G.; conducted data analysis and interpretation: K.R.A., E.F.C., T.E.B., T.S., R.A-D., S.S.H., E.M.T.; drafted manuscript: K.R.A.; Commented, edited, approved manuscript: all.

## Acknowledgments

We thank the following people and organizations assistance with sample collection: Dave Casebolt and Miranda Holt from the University of Idaho Sheep Center; Brandon Munk and Erin Schaeffer from the CA Department of Fish and Wildlife; Brandi Felts, Jon Jenks, Laura McHale from South Dakota State University; Hank Edwards from Wyoming Game and Fish Department Wildlife Health Laboratory; Kerry Sondgeroth, Kevin Monteith, and Brittany Lyn Wagler from the University of Wyoming; Bob Gerlach from the Office of the State Veterinarian, Alaska; Shari Willmott and Helen Schwantje from British Columbia Wildlife Health Program; Kerry Mower from New Mexico Department of Game and Fish; Peri Wolff, Matt Jeffress, Mike Cox Nevada Department of Wildlife; Jace Taylor, Utah Division of Wildlife Resources; R. Scott Larsen from the Denver Zoological Foundation; Emily Almberg from Montana Fish, Wildlife & Parks. We thank Digpal S. Gour, Brandon Larsen, and Kara Robbins for assistance with laboratory work. Mark McGuire assisted with facilitating this project through the University of Idaho College of Agriculture and Life Sciences.

