## Supplemental_Text for "Comparative genomic analysis identifies potential adaptive variation and virulence factors in *Mycoplasma ovipneumoniae*"

### Supplemental Materials

### Methods

*Genome assembly*

For samples with Oxford Nanopore Technologies (ONT) sequence data, raw ONT sequence reads were demultiplexed by barcode and base-called using Guppy v.4.2.2 (Oxford Nanopore Technologies Ltd.). Assembly was performed using the Trycycler pipeline (https://github.com/rrwick/Trycycler), which incorporates methods for identifying and removing potential contamination from assemblies. First, we removed short reads (less than 500bp) and low-quality reads (the 5% of reads with the lowest quality scores) for each sample. Next, we subsampled the filtered reads 18 times for each sample, and used these subsampled reads to perform six assemblies with each of three different programs: Flye [1], Miniasm+Minipolish [2] (https://github.com/rrwick/Minipolish), and Raven [3]. All assembled contigs were clustered by sequence similarity for each sample, and dendrograms of contigs were visualized using Dendroscope [4] to identify and select clusters containing contigs of the expected assembly length (approximately 1MB for *M. ovipneumoniae*) and read depth. The contigs within each selected cluster were then "reconciled" through a process that involves circularizing contigs and removing contigs that could not be circularized or had lengths or sequence compositions inconsistent with the other contigs. Multiple sequence alignment was then performed with reconciled contigs, and filtered sequence reads that assigned to the reconciled contigs were identified. Next, a consensus sequence was generated for the assembly, and assemblies were polished with the ONT reads that assigned to the reconciled contigs using Medaka (Oxford Nanopore Technologies Ltd.). When Illumina sequence data were available for the same sample, assemblies were further polished with the cleaned Illumina sequence reads by performing two rounds of polishing with Pilon [5]. Depth of coverage was calculated using SAMtools v.1.5 [6] after mapping cleaned reads to the assembly using Miniasm for ONT reads, and BWA v0.7.17 [7] for Illumina reads.

For samples with only Illumina sequence data and no ONT sequence data (due to insufficient amounts of genomic DNA after culture enrichment, as described in the main text), cleaned Illumina reads were assembled de novo using SPAdes v3.13.1 [8], using the parameters --careful and -k 33,77,127. This method was also used to assemble genomes for the raw Illumina sequence reads downloaded from NCBI for six samples collected in France in a previous study, as described in the main text [9]. For each sample, cleaned reads were mapped back to the assembly using BWA and depth of coverage was calculated using SAMtools.

### Results

*Genome assembly statistics*

Of the 74 genome assemblies we generated for this project, 46 had both ONT and Illumina data, and 28 had only Illumina data (Table S1). Of the assemblies with both ONT and Illumina data, all except one was circularized ("closed"); for these, the mean depth for the ONT data ranged from 29.07 to 1,847.23 (mean = 448.96) and for the Illumina data ranged from 44.43 to 411.15 (mean = 201.59). For the assemblies with only Illumina data, N50 ranged from 12,857 to 586,790 (mean = 83,586.39); the number of contigs ranged from 49 to 647 (mean = 211.93); and mean depth ranged from 66.88 to 494.09 (mean = 161.27). Of the 25 assemblies we obtained from NCBI, two were generated by PacBio Sequel II sequencing, three by PacBio RS sequencing, one by Roche 454 sequencing, and the rest were generated by Illumina sequencing. For the PacBio assemblies, four were circularized, and the fifth had two contigs and N50 = 678,446; mean depths ranged from 377.46 to 2,350. For the Roche 454 assembly, N50 = 47,128, number of contigs = 49, and mean depth = 60. For the Illumina assemblies, the N50 ranged from 16,307 to 571,788 (mean = 80,139.26); the number of contigs ranged from 13 to 143 (mean = 81.84); and mean depth ranged from 31 to 2,857 (mean = 1,012.00). Across all assemblies in our final dataset, the assembly lengths ranged from 799,992 to 1,365,101 bp (mean = 1,083,003)(Fig. 5a, Table S1).

*CRISPR-Cas systems*

We identified three genes associated with CRISPR-Cas systems (*cas1*, *cas2*, *cas9*) in almost all genomes (Table S2, S12). *cas1* and *cas2* were absent from only five genomes, including all three caribou genomes and the two OCAHG genomes. *cas9* was absent from three genomes from the sheep clade, including the genome sampled in Brazil, one sampled at a USA farm, and one of the type strains from Australia (OARBZ_USPBR2017, OARF1_109, OARAU_ATCC2). However,

11 additional genomes had only partial *cas9* sequences (i.e. covering <50% of the consensus sequence from the multi-sample nucleotide alignment), possibly indicating a nonfunctional gene. These 11 samples included the 3 caribou genomes, the two OCAHG genomes, one of the type strains (OARAU_ATCC1), and five additional bighorn sheep genomes (all from different populations). In addition, four genomes had two copies of *cas9* (OARF4_W260, OARFR_F11630, OCABB_370144, OCAFR_370127). CRISPR repeat regions were present in 95 samples, and absent in 4 samples (OARAU_ATCC2, OCAHC_KL3363, OCAHG_370152, OCAHG_370153) (Table S12, S13). The number of repeats ranged from 4 to 188, with a mean of 58. The three caribou samples had the lowest numbers of repeats (two samples had 4 repeats, one had 5 repeats; the next highest number of repeats was 9 for sample OCALO_20LO68). Thus, our analyses indicate that most isolates have functional CRISPR-Cas systems, with the exception of the three caribou samples and the two bighorn sheep genomes from HG, and possibly nine additional samples (the two type strains, two additional domestic sheep, and five additional bighorn sheep samples). Notably, gene absence (or partial presence) could result from genome assembly errors, culturing, or divergence from the reference database.

*MLST analyses*

BLAST+ identified two assemblies (CHIBN_274 and OARBN_150) that had incomplete sequences of the 16S marker, and these sequences were removed from subsequent analyses. This analysis also identified two assemblies (CHIAS_Crony and OCABH_190781) that had two copies of the gyrB marker. For CHIAS_Crony, the two copies had identical sequences, and so we retained one copy of gyrB for subsequent analyses for that sample. For OCABH_190781, the two copies had different sequences, and so we completely removed that sample from subsequent analyses. After sequence alignment, the lengths of each MLST were 394bp for gyrB, 355 bp for IGS (after removing a region with a large number of indels), 343bp for 16S, and 562bp for rpoB.

### Supplemental Figures

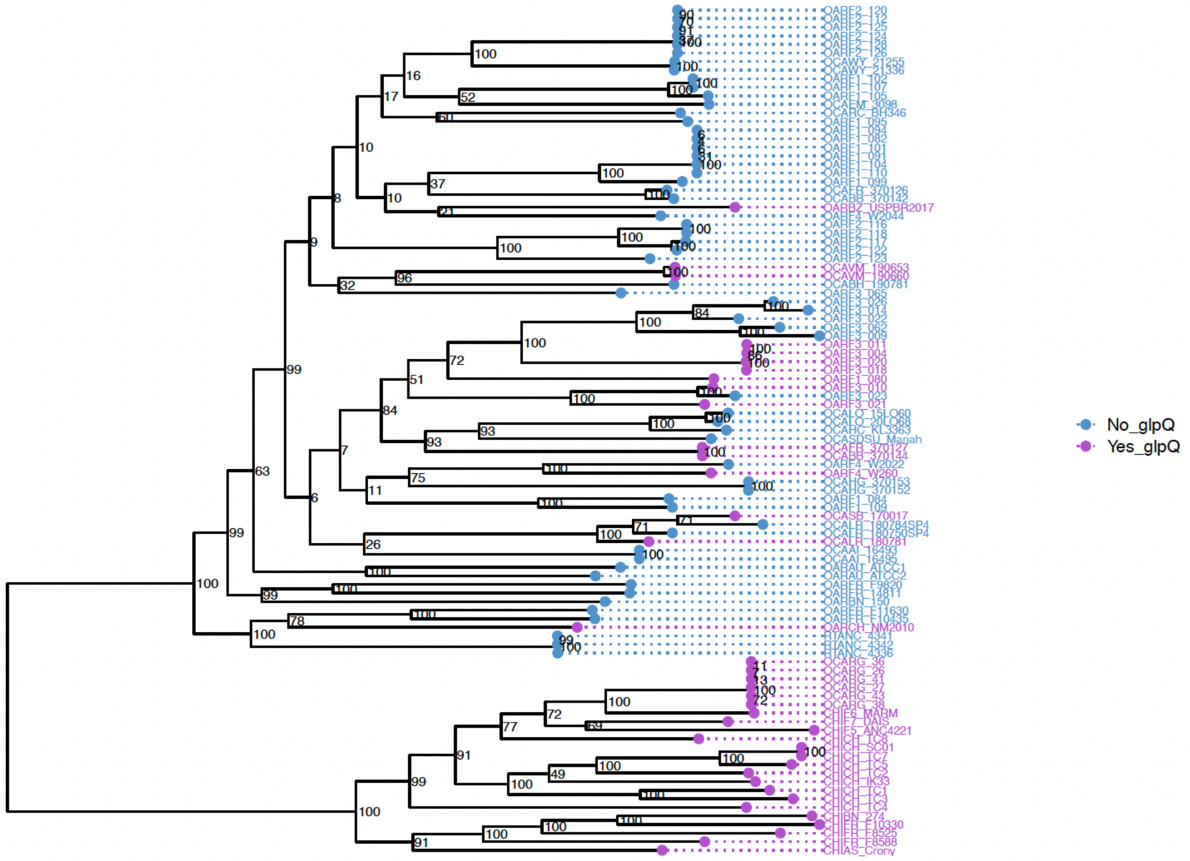


Figure S1. Maximum likelihood phylogeny of core genome sequences for 99 *Mycoplasma ovipneumoniae* assemblies, with colors representing the presence ("Yes_glpQ") or absence ("No_glpQ") of the gene *glpQ* in the genome assembly. The presence of this gene was significantly associated with the goat clade.

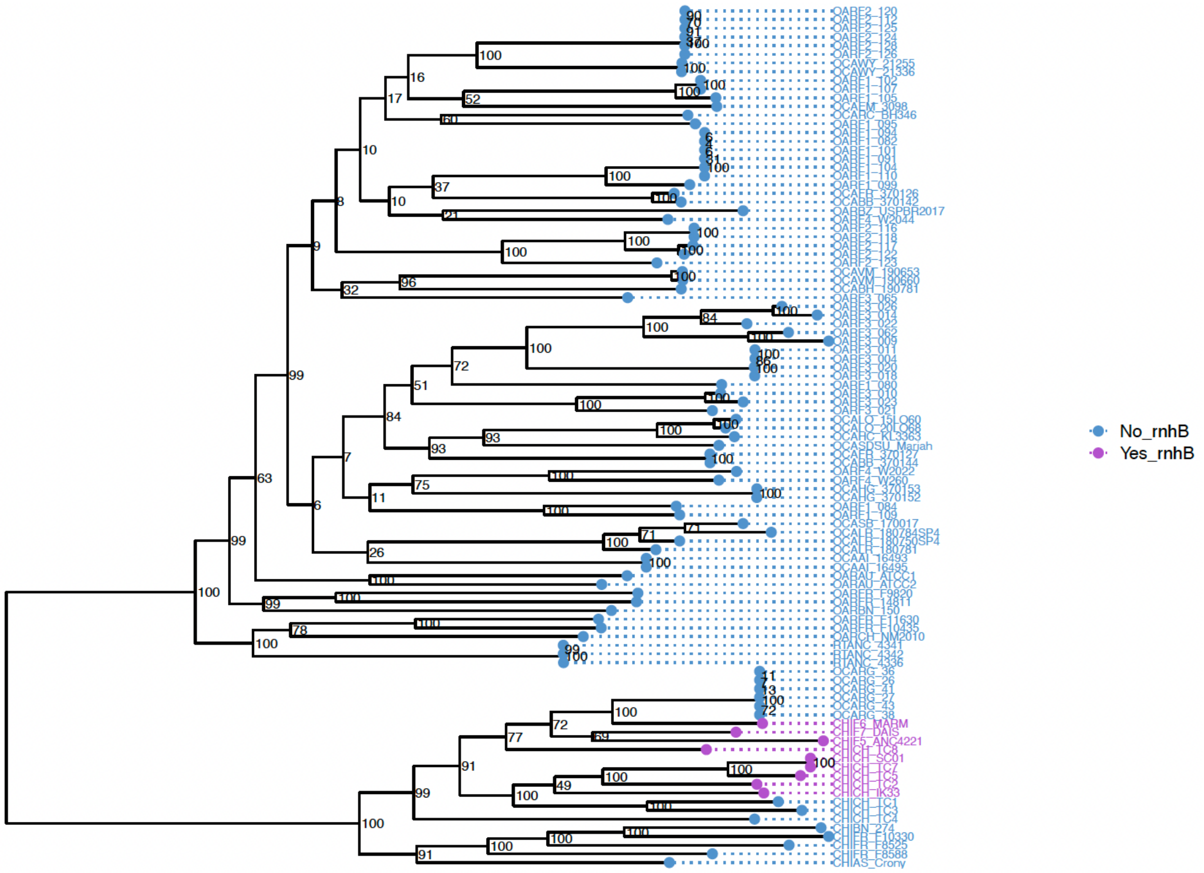


Figure S2. Maximum likelihood phylogeny of core genome sequences for 99 *Mycoplasma ovipneumoniae* assemblies, with colors representing the presence (“Yes_rnhB”) or absence (“No_rnhB”) of the gene *rnhB* in the genome assembly. The presence of this gene was significantly associated with the goat clade.

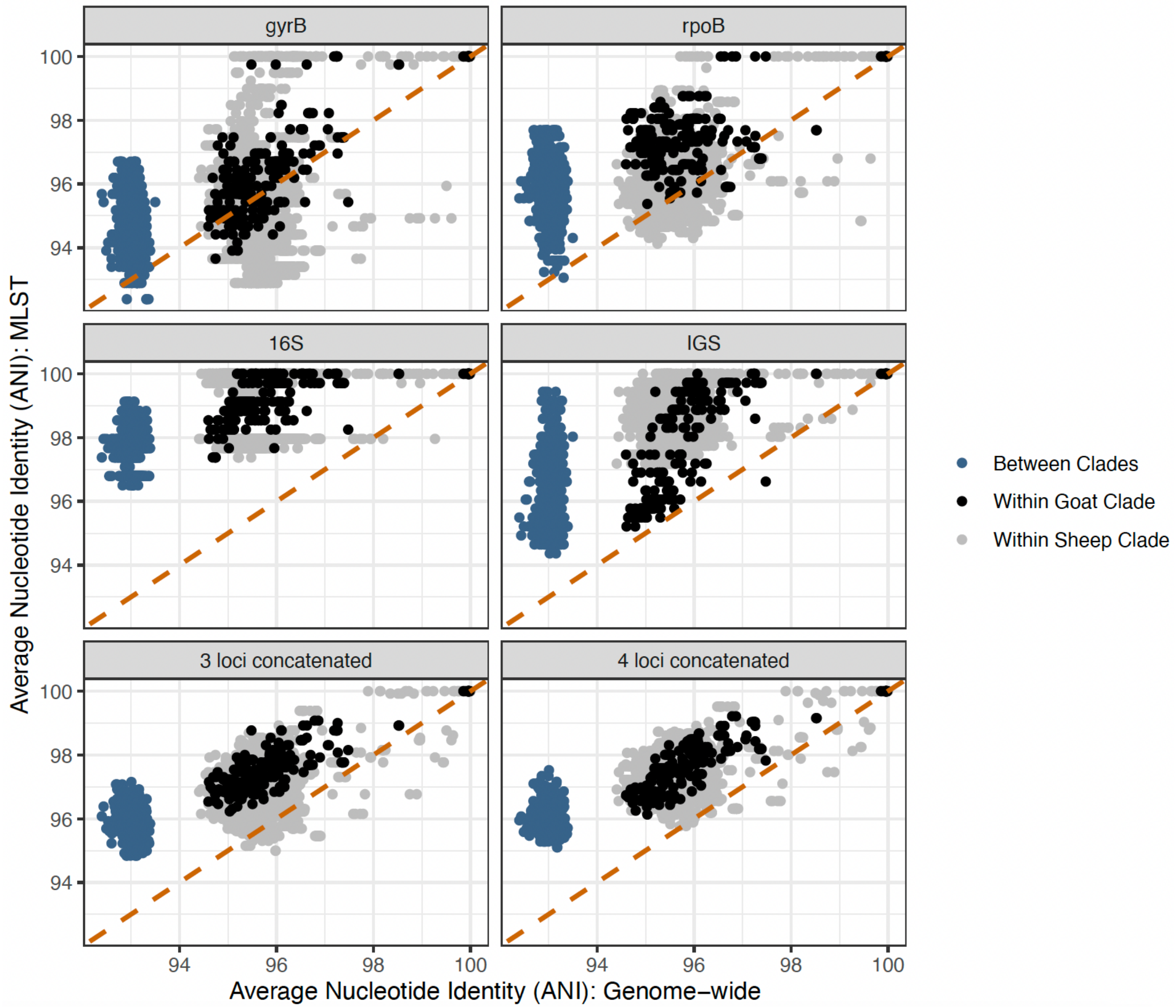


Figure S3. Comparison of Average Nucleotide Identity between each pair of samples calculated using sequences from multi-locus sequence typing (MLST) markers, or from genome assemblies. “3 loci concatenated” = results for analyses using concatenated gyrB, rpoB, and 16S sequences; “4 loci concatenated” = results for analyses using concatenated sequences for all four markers. Orange dashed line represents a hypothetical relationship in which ANI values calculated with MLST markers are identical to those calculated with genome assemblies.

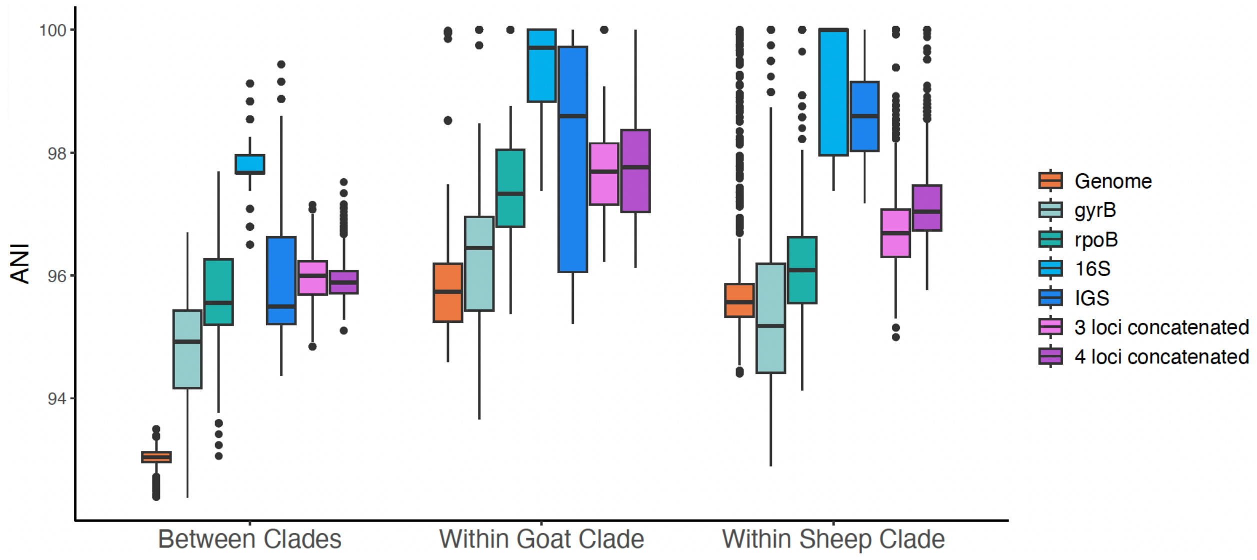


Figure S4. Boxplots of Average Nucleotide Identity between each pair of samples calculated using sequences from multi-locus sequence typing (MLST) markers, or from genome assemblies. “3 loci concatenated” = results for analyses using concatenated gyrB, rpoB, and 16S sequences; “4 loci concatenated” = results for analyses using concatenated sequences for all four markers.

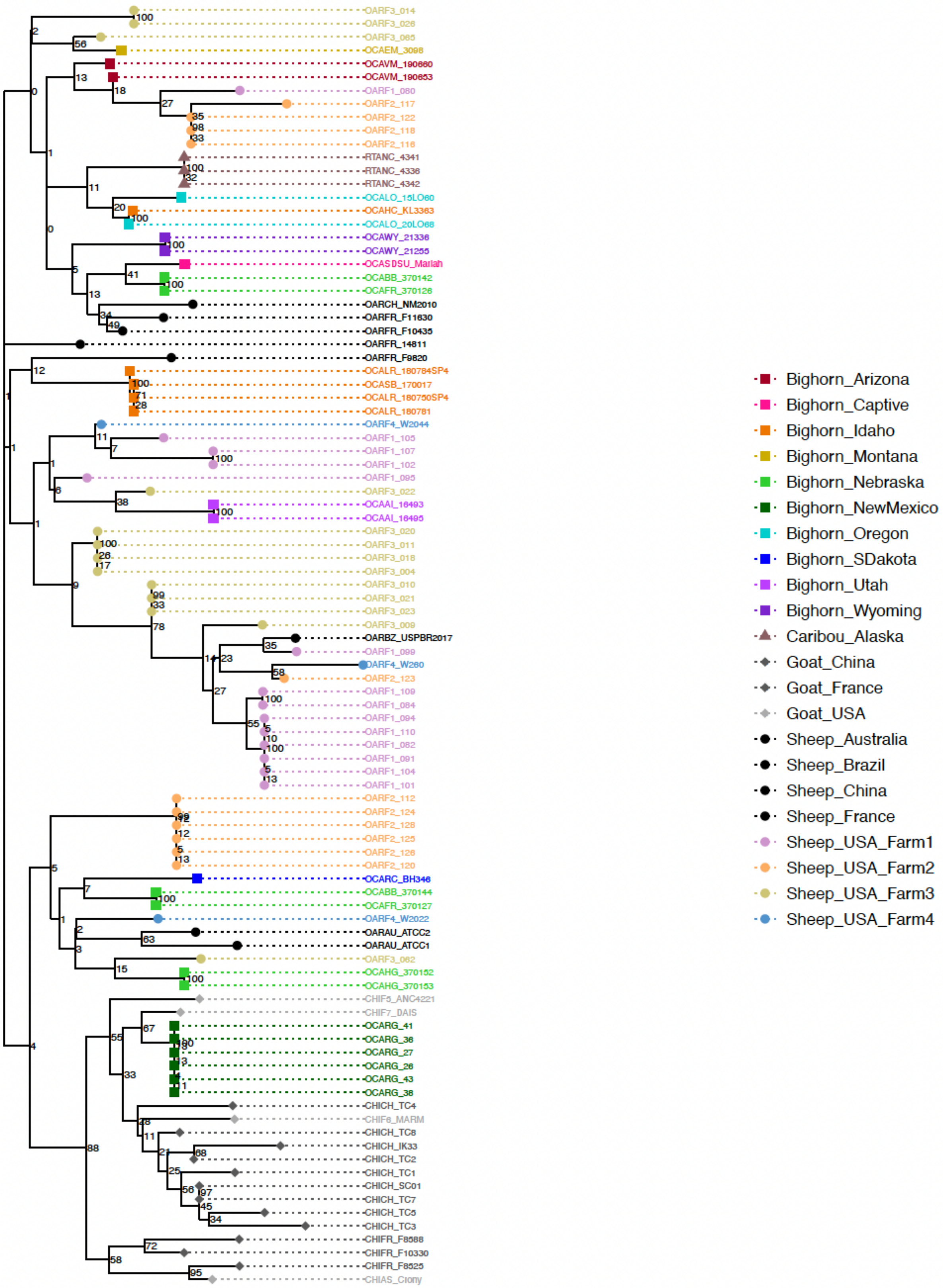


Figure S5. Maximum likelihood phylogeny for *Mycoplasma ovipneumoniae* generated using sequences from three concatenated MLST markers (gyrB, rpoB, 16S).

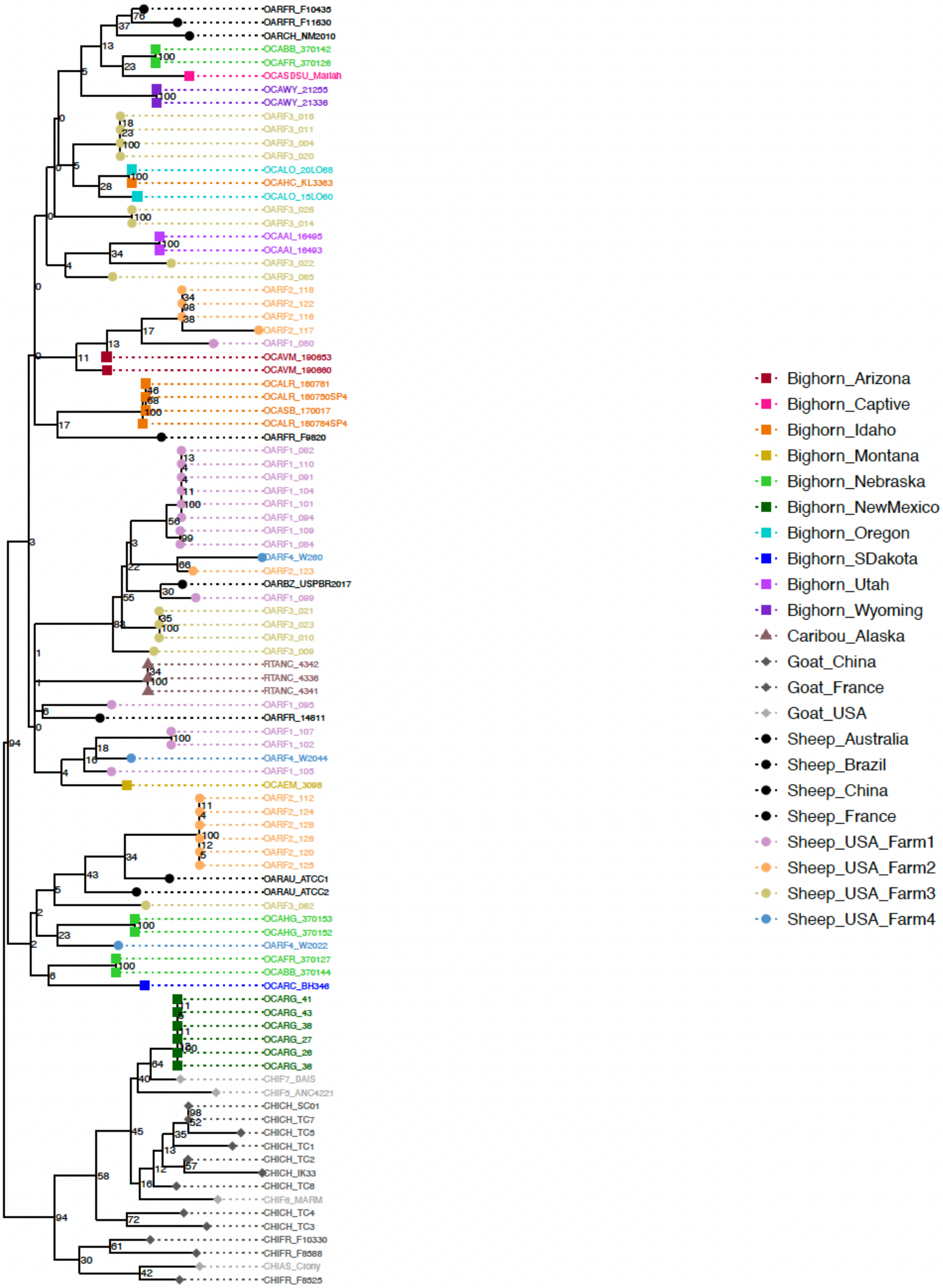


Figure S6. Maximum likelihood phylogeny for *Mycoplasma ovipneumoniae* generated using sequences from four concatenated MLST markers (gyrB, rpoB, 16S, IGS).
